# An organoid co-culture model for probing systemic anti-tumor immunity in lung cancer

**DOI:** 10.1101/2024.06.04.597327

**Authors:** Kaiyi Li, Chang Liu, Chao Li, Ting Zhang, Tian Zhao, Dong Zhang, Hainan Wu, Yuhan Liu, Shuai Wang, Yingshun Yang, Baobao Lin, Wenyan Wang, Jun Wang, Xizhao Sui, Xiaofang Chen, Peng Liu

## Abstract

Deciphering the interactions between tumor micro- and systemic immune macro-environment is essential for developing more effective cancer diagnosis and treatment strategies. Here, we established a gel-liquid interface (GLI) co-culture of lung cancer organoids (LCOs) and paired peripheral blood mononuclear cells (PBMCs), featuring with enhanced interactions of immune cells and tumor organoids, to mimic the *in vivo* systemic anti-tumor immunity induced by immune checkpoint inhibitors (ICI). The co-culture model recapitulates the *in vivo* ICI-induced T cell recruitment and subsequent tumor regression, predicting the clinical results precisely. We demonstrated that circulating tumor-reactive T cells, which are effector memory-like with high expression levels of *GNLY, CD44* and *CD9,* can serve as an indicator of the immunotherapy efficacy. Interestingly, enhanced inflammatory signaling in blood T cells is accompanied with prompted exhaustion and compromised anti-tumor function, when encountering with organoids. Our findings suggest that the GLI co-culture can be used for developing diagnostic strategies for precision immunotherapies as well as understanding the underlying mechanisms.

**Summary figure:** 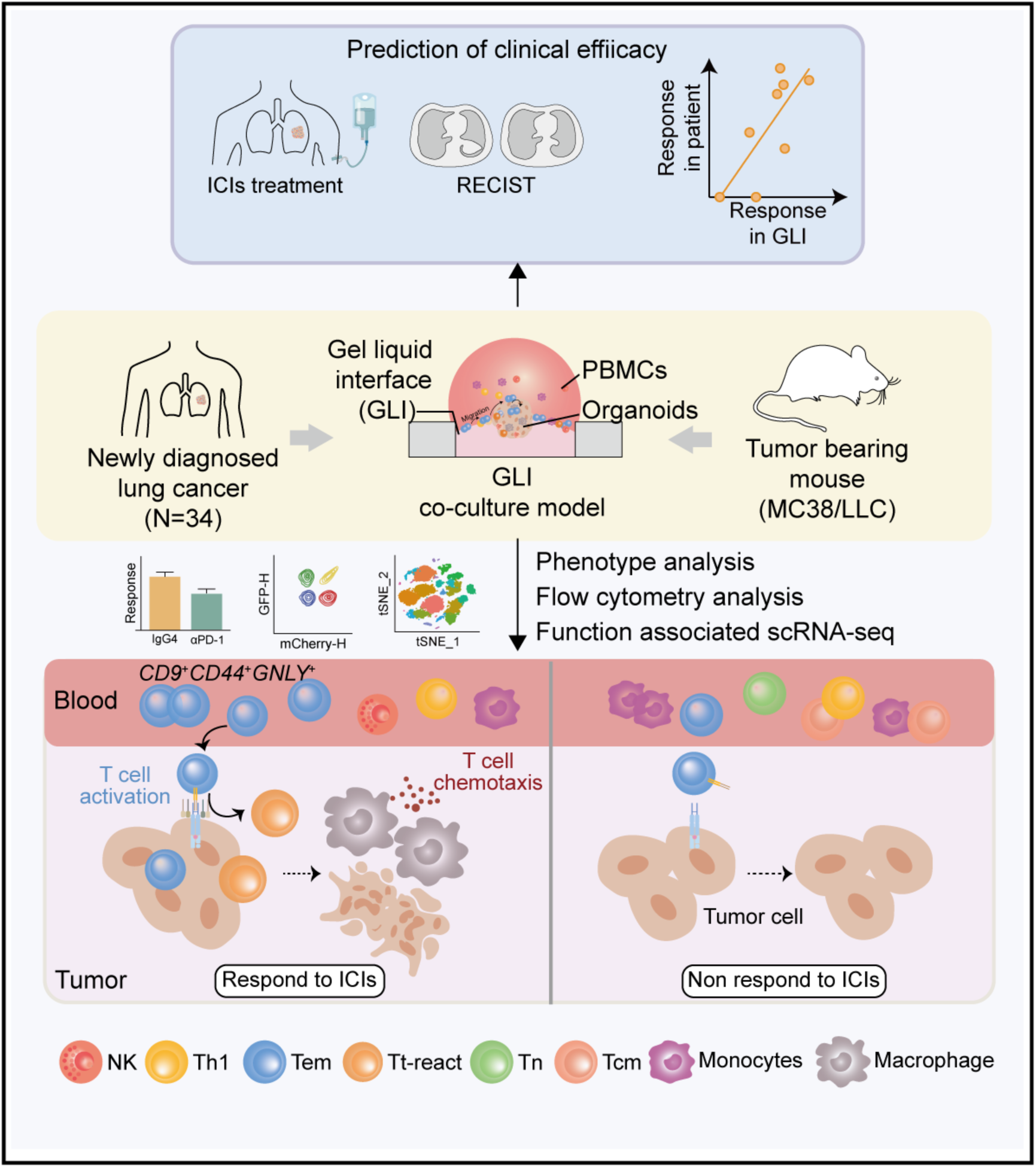

## Main

Lung cancer is one of the most prevailing malignancies in both genders worldwide, accounting for an annual incidence of 2,206,771 cases and 1,796,144 fatalities.^1,2^ Over the past decades, immune checkpoint inhibitors (ICI) have revolutionized the landscape of cancer treatment and been approved by FDA for use in diverse cancer types, particularly improving the clinical outcomes of advanced-stage lung cancer.^3,4^ However, the remarkable responses to ICI-based immunotherapies are limited to a minority of patients. A series of characteristics of tumors have been utilized as predictive biomarkers for ICI responses, such as PD-L1 expression of tumor cells, T cell infiltration into tumor tissues, tumor mutation burden, etc.^5^ Unfortunately, none of these indices have been proven trustworthy at all time, due mainly to the complex interactions between tumor and the macro-environment of the host body. The anti-tumor immunity is coordinated across tissues, and the localized anti-tumor immune response cannot persist without continuous communication with the peripheral immune cells. Actually, recent observations have shown that post-treatment T cells were significantly enriched for novel clonotypes which may be recruited from extratumoral sites (i.e., peripheral blood and draining lymph nodes).^6–8^ Additionally, the identification of circulating neoantigen-specific CD8^+^ T cells in the blood substantiated the involvement of circulating T cells in anti-tumor immune surveillance and treatment.^9^ Yet, due to the lack of efficient tracking tools, how cells in tumor circulating systems dynamically respond to ICI and interact with local tumor microenvironment (TME) remains elusive.

Advancing techniques, such as single-cell sequencing,^8,10^ multiplex high-resoultion imaging techniques,^11^ and spatial transcriptomics,^12^ provide high-resolution cell atlases in TMEs and circulating systems across cancer types, identifying a spectrum of functional and dysfunctional states of T cells. Unfortunately, due to the inherent challenge of acquiring longitudinal tumor samples from patients, only snapshots at certain time points, typically before or after treatments, can be obtained,^13,14^ making it impossible to directly track T cell differentiation, proliferation, and tumor killing over the time. While window-of-opportunity experiments offer some insights into T cell fate, they are limited in temporal scope,^6,7,14^ underscoring the critical need for models to comprehensively elucidate the dynamics of anti-tumor immunity. Recent advances include a methodology demonstrating the tracking of tumor-specific CD8^+^ T cell populations in a standardized and temporally calibrated *in vivo* mouse model.^15^ Nevertheless, the lack of patient-specific diversity compromises the clinical revelence of this approach. Conversely, an easily accessible *ex vivo* model mimicing the systemic immune response accurately offers an alternative strategy for dissecting the dynamics of the anti-tumor immunity during the course of ICI treatment.

The arising of tumor organoid model has bridged the gap between the basic research and the clinical practice, demonstrating the capability of predicting patient-specific outcomes under chemo- and targeted-therapies.^16,17^ The application of tumor organoid models has also been extended to the area of anti-tumor immunity in recent years. Patient-derived organoids (PDOs), such as air-liquid interface model, comprising tumor epithelium and stroma with infiltrated immune cells,^18^ have facilitated the assessment of immunotherapy.^19^ More recently, the reconstituted models consisting of PDOs and exogenous immune cells have been reported, demonstrating expanding of tumor-specific CD8^+^ T cells^20^ and CAR-NK-mediated cytotoxicity toward tumor organoids.^21^ However, whether such models can mimic the interactions between local tumor and systemic immunity, and how they can be used to dissect the dynamic immune responses induced by ICI are still unclear.

Here, we established a gel-liquid interface (GLI) co-culture model, where patient-derived lung cancer organoid (LCOs) were positioned at the interface of Matrigel and culture medium for efficient interactions with peripheral blood mononuclear cells (PBMCs) from the same patient. Subsequently, we demonstrated the clinical efficacy of immunotherapies could be accurately reflectly by the GLI co-culture model. Based on the function-associated single-cell RNA sequencing (FascRNA-seq) platform,^22^ we employed multi-omics approaches to identify the molecular characteristics of circulating T cells related to patient responses, as well as depicting the dynamic process of T cell infiltration and cytotoxic function. Finally, we dissected the responding patterns of circulating T cells correlated to the features of the tumor tissues. These findings deepen our understanding on mechanisms related to ICI responses and offer insights into systemic immune response in lung cancer (Fig. 1a).

**Fig. 1.**
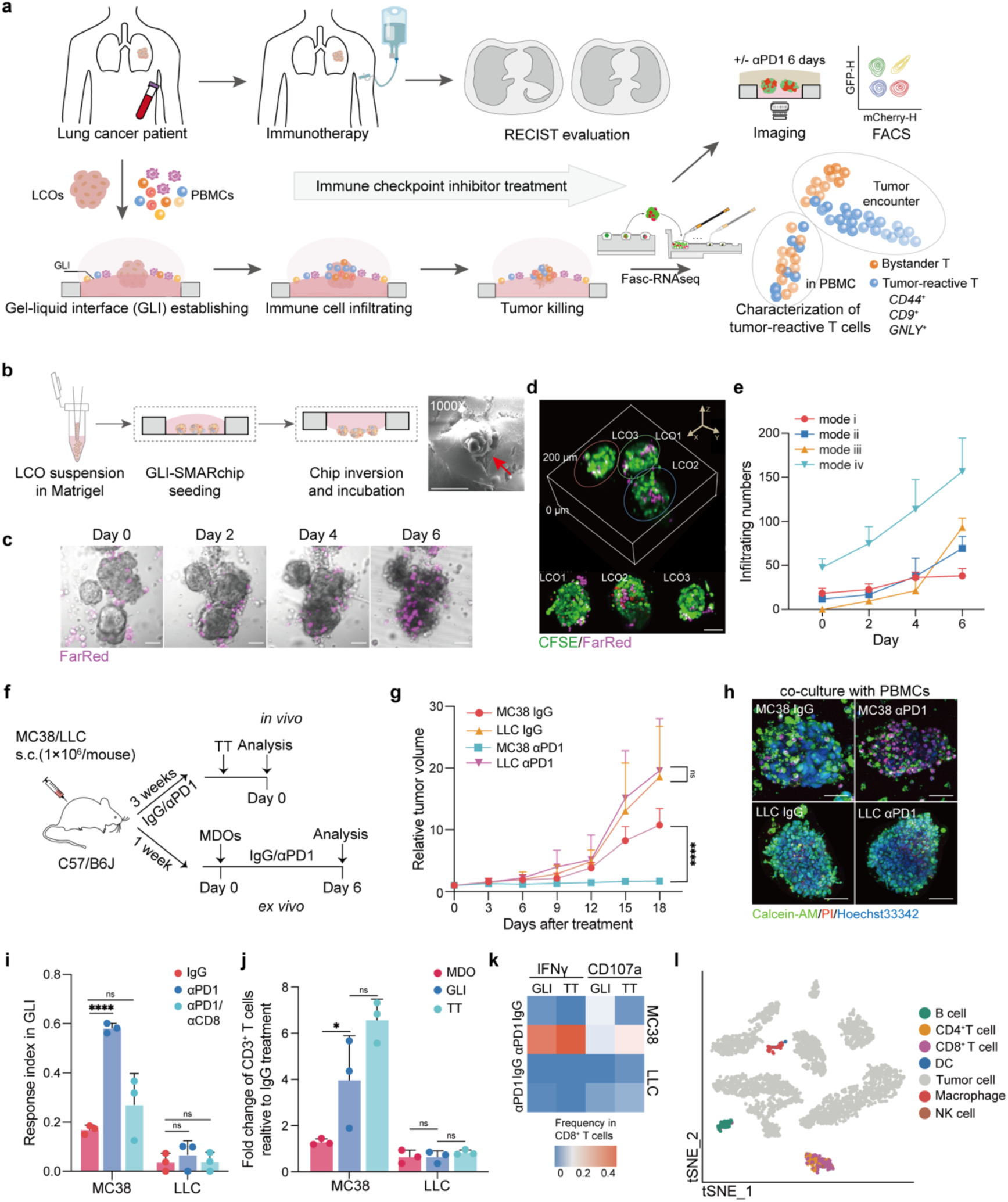
Establishment and validation of the GLI co-culture model. **a,** Schematic diagram illustrating the analysis of ICI-induced dynamic immune responses in the GLI co-culture and characterization of tumor-reactive T cells. **b,** Key steps of GLI co-culturing (left) and scanning electron microscopy image showing an organoid at the GLI (right), the red arrow represents the LCO. Scale bar: 50 μm. **c,** Representative images depicting PBMCs migrating and infiltrating into LCOs in GLI co-culturing. PBMCs were labeled with FarRed (purple). Scale bars: 50 μm. **d,** Representative 3D images of LCOs with infiltrated PBMCs in GLI co-culturing. LCOs and PBMCs were labeled with CFSE (green) and FarRed (purple), respectively. Scale bar: 50 μm. **e,** Comparison of the number of PBMCs infiltrated into LCOs over time under different co-culture modes (n=10). Mode i: mixing LCOs with PBMCs in the medium; mode ii: mixing LCOs with PBMCs in Matrigel; mode iii: embedding LCOs alone in Matrigel and model iv was the GLI co-culture. **f,** Experimental design of animal experiments for validation of the GLI co-culture. **g,** Relative tumor volumes of MC38 and LLC following αPD1 treatment versus isotype control IgG (n = 6). **h,** Fluorescence microscopy showing the viability of MC38 and LLC GLI co-cultures on day 6 under αPD1 or IgG treatment. Live cells and dead cells were labeled with Calcein-AM (green) and PI (red), respectively. Scale bars: 50 μm. **i,** Response index (Ri) of MC38- and LLC-derived GLI models following IgG or αPD1 treatment (n = 3). **j,** FACS analysis of the fold change of CD3^+^ T cells in MC38- and LLC-derived MDO mono-cultures, GLI co-cultures and TTs after αPD1 treatment relative to IgG (n=3). **k,** Heatmap showing the IFNγ^+^CD8^+^ T cell and CD107a^+^CD8^+^ T cell frequencies in MC38- and LLC-derived GLI models and TTs following IgG or αPD1 treatment. **l,** t-SNE visualization of cells from MDO and GLI models labeled by cell types. (\**P*<0.05, \*\*\*\**P*<0.0001, ns: no significance by unpaired student’s *t* tests)

## Results

### Establishment of GLI co-culture model

To enhance the interactions between the tumor organoids and immune cells while maintaining their viabilities, we established a brand-new gel-liquid-interface co-culture model on a superhydrophobic microwell array chip (GLI-SMARchip) developed previously by our group,^23^ which contains an array of 1.5-μL microwells with a thin glass bottom for confocal imaging and a layer of superhydrophobic polymers on the upper side for microwell isolation (Extended Data Fig. 1a,b). In the GLI co-culture model, LCOs were positioned mostly below the surface of Matrigel in the microwells and PBMCs were cultured on the surface of the gel (Fig. 1a,b). To achieve this cell arrangement, LCOs mixed with Matrigel were first seeded into the microwells, and then the chip was kept upside down on ice to induce the sedimentation of LCOs towards the Matrigel surface under gravity (Fig. 1b and Extended Data Fig. 1c). Once the gel was solidified, the chip was inverted and PBMCs in T cell culture medium were pipetted onto the gel surface to form the GLI co-culture (Extended Data Fig. 1c). We optimized the sediment time to 10-minute and the Matrigel concentration to 80% so that the LCOs can stably remain right on the edge for up to 6 days (Extended Data Fig. 1d,e). Two LCO lines (LCO 4853 and LCO 4841) established previously cultured in the GLI for 7 days demonstrated the similar viabilities and growth rates as those in Matrigel, better than those suspended in the culture medium without gel (Extended Data Fig. 2a-c). The viability of the organoids cultured in the GLI was assessed by Calcein-AM/PI staining followed by confocal imaging and further confirmed by flow cytometry (Extended Data Fig. 2g). The maintenance of organoid viability in GLI with the T cell culture medium ensured the proper engagement of immune cells with the tumor.

Unlike the conventional hanging-drop method in which the droplet usually had a large curvature,^24^ a relative flat gel surface could be stably formed within the 1.2-mm-diameter microwells even in the upside-down position due to the surrounding superhydrophobic polymers on the GLI-SMARchip. Such a flat gel surface enabled the free migration of immune cells towards organoids, thereby improving the infiltration (Fig. 1c,d). We compared the infiltration numbers of the immune cells in different co-culture modes: i) mixing organoids with PBMCs in medium, ii) mixing organoids with PBMCs in Matrigel, iii) embedding organoids alone in Matrigel, and iv) the GLI. By testing a pair of PBMCs and organoids obtained from a lung cancer patient, the GLI model demonstrated the most effective infiltration of the PBMCs as well as the killing of tumor cells under the treatment of αPD1 (Fig. 1e and Extended Data Fig. 2d-f). Additionally, the structure of the GLI allowed us to easily remove the PBMCs not engaged with LCOs and then to harvest the LCOs for subsequent single-cell sequencing and flow cytometry analysis.

### GLI models derived from syngeneic mouse tumors mimic *in vivo* immune responses to ICI

To evaluate whether the GLI models can mimic the *in vivo* anti-tumor immune response to ICI, mouse-derived organoids (MDOs) were first generated from MC38 and LLC mouse tumors following the protocol that developed previously by our group (Extended Data Fig. 3a).^25^ We have proved that considerable amounts of lymphocytes could be preserved in the MDOs (Extended Data Fig. 3b).^22^ The MDOs were then co-cultured with PBMCs in the GLI mode followed by the treatment with αPD1 or control IgG (Fig. 1f). Following the αPD1 treatment, we observed that the tumor growth was impeded in MC38 but not LLC mice (Fig. 1g), in aggrement with previously reported ICI sensitive (MC38) and resistant (LLC) phenotypes.^26^ The *ex vivo* GLI models demonstrated the similar responses as the *in vivo* tumors where significant cell death was observed in the αPD1-treated MC38 but not the LLC models (Fig. 1h,i). Furthermore, the response to αPD1 was compromised by the addition of αCD8 (Fig. 1i), indicating the critical role of CD8^+^ T cells for αPD1-induced anti-tumor immunity.

Since αPD1 was also reported to enhance immune cell infiltration,^27^ we examined the frequencies of immune cells infiltrated into both the tumor tissues (TTs) and the MDOs in the GLI co-cultures using fluorescence-activated cell sorting (FACS). The infiltration of CD3^+^ T cells was enhanced significantly in both the MC38-derived GLI models and the TTs under αPD1, whereas the LLC-derived GLI models and TTs exhibited a decline in T cell frequencies (Fig. 1j and Extended Data Fig. 3c). Interestingly, the T cell frequencies in MDOs cultured without PBMCs did not show such significant changes (Fig. 1j and Extended Data Fig. 3c), indicating the migration of T cells from PBMCs into the MDOs in the GLI co-culture (Extended Data Fig. 3d,e).

Consistently, the IFNγ and CD107a productions, the indicators of T cell activation and cytotoxic function, were prominently upregulated in CD8^+^ T cells collected from MC38-derived TTs and GLI models under the αPD1 treatment, while LLCs showed no significant changes (Fig. 1k). The same trends were seen at the mRNA levels of *Ifng, Prf1*, and *Gzmb* (Extended Data Fig. 3f). FascRNA-seq was employed to further analyze the immune cells infiltrated into MDOs and multiple types of immune cells, including macrophages, natural killer cells, and dendritic cells were detected in MDOs (Fig. 1l), consistent with our previous report.^22^ Consistent with the FACS and qRT-PCR data, the genes associated with T cell activation and cytoxicity were significantly upregulated in the MC38 GLI co-culture with αPD1, indicating a more activated and effector-like state (Extended Data Fig. 3g). These results demonstrated that the GLI co-culture recapitulates the ICI-induced *in vivo* immune responses in mouse tumors.

### 1GLI co-culture model recapitulates patient response to immunotherapy

Next, we generated LCOs and established GLI co-cultures from 34 treatment naïve lung cancer samples, including 25 adenocarcinomas (ACs), 5 squamous cell carcinomas (SCCs) and 3 small cell lung cancers (SCLCs), information of patients are listed in Table 1. The cytotoxic effect induced by nivolumab alone (αPD1) or nivolumab plus chemotherapy (αPD1/Chemo), as indicated by the response index (Ri, the ratio of dead tumor cells and total tumor cells), were evaluated for all the samples (Fig. 2a, Table 2). As expected, Ri was significantly elevated by αPD1 and further increased by αPD1/Chemo (Fig. 2b), consistent with the results in previously reported clinical trials.^28^ Based on the range of Ri in the IgG4 group (0.161 ± 0.084), we determined Ri = 0.245 (mean ± SD) as the threshold to distinguish immunotherapy responder (R) or non-responder (NR) samples (Fig. 2b). No significant difference of Ri was observed among the subtypes of lung cancer (Fig. 2c and Extended Data Fig. 4a), while the responses to both αPD1 and αPD1/chemo were compromised in the stage IV samples (Fig. 2d and Extended Data Fig. 4b), echoing advanced immune escape in late-stage patients.^29^ Interestingly, Ri of GLI models were significantly higher than that in the LCO alone (Fig. 2e and Extended Data Fig. 4c), suggesting the anti-tumor roles played by PBMCs.

**Fig. 2.**
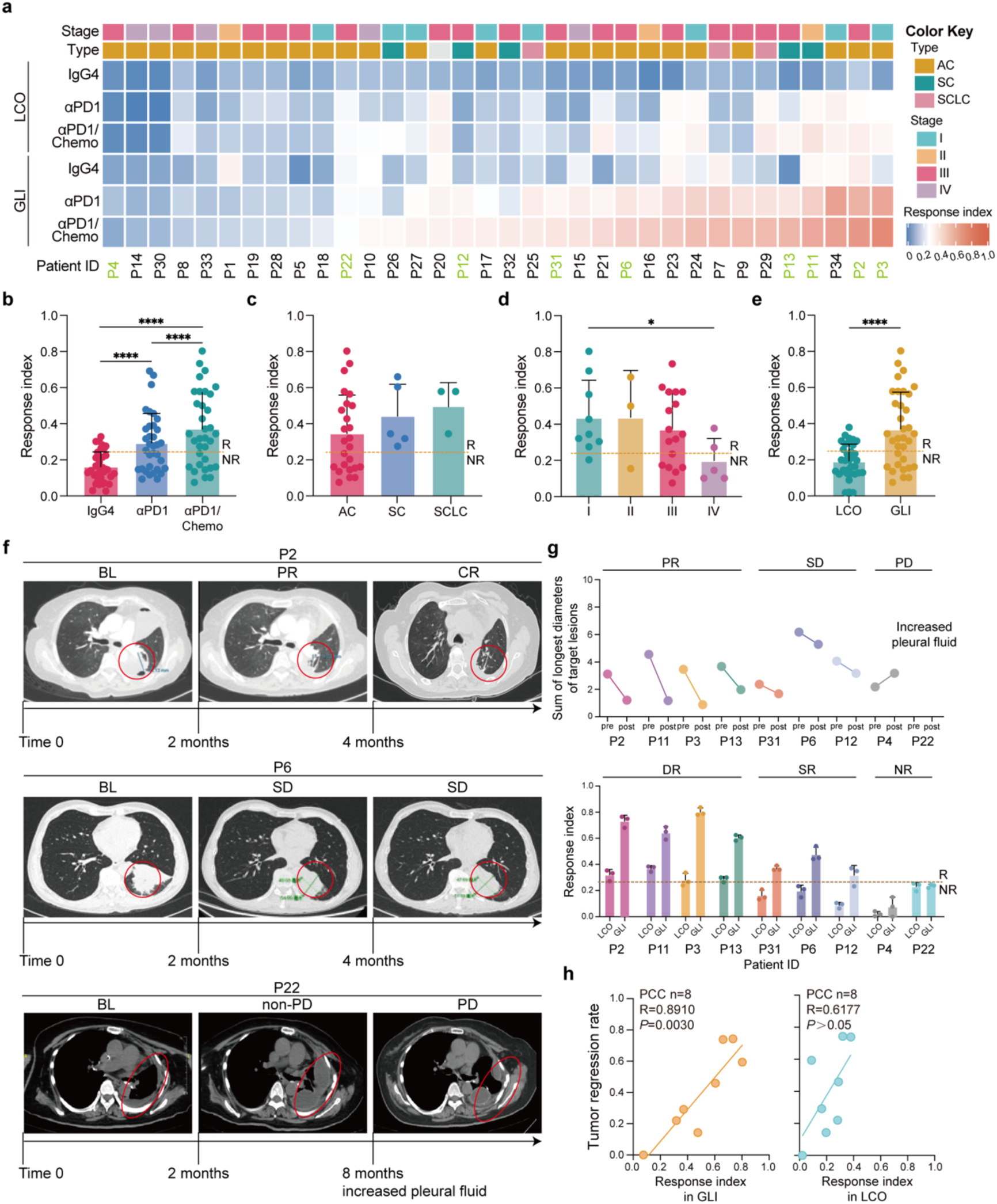
Response of GLI co-culture represents the clinical results of immunotherapy. **a,** Ri of the LCO mono- and GLI co-cultures under different treatment conditions including IgG4, αPD1 or αPD1/chemo. The green codes indicate patients receiving immunotherapy *in clinic*. **b,** Bar plots comparing Ri of GLI models under different treatment conditions (n=34, \*\*\*\**P* <0.0001 by paired student’s *t* tests). **c,d,** Bar plots comparing Ri of GLI models derived from patients with different subtypes (**c**) and stages (**d**) treated with αPD1/chemo, note the significant decrease of Ri for the stage IV group (\**P*<0.05 by unpaired student’s *t* tests). **e,** Bar plots comparing Ri of LCO mono- and GLI co-cultures following αPD1/chemo treatment (n=34, \*\*\*\**P*<0.0001 by paired student’s *t* tests). **f,** CT scan images showing the temporal evolution of lesions in patients P2, P6 and P22 undergoing immunotherapy and the RECIST evaluation results. The red circles indicate the primary tumor (top and middle) and the pleural fluid (bottom). **g,** The sum of longest diameters (SLD) of target lesions pre- and post-immunotherapy (top), and Ri of the corresponding LCO and GLI models (bottom). **h,** PCC analysis between the tumor regression rate (the decreasing ratio of SLD for target lesions) and Ri of the GLI co-cultures (left), LCO mono-cultures (right).

Nine of the 34 patients received the αPD1-based therapies *in clinic*, where 4 patients had partial response (PR), 3 had stable disease (SD) and the other 2 had progressed disease (PD) according to RECIST^30^ (Fig. 2f,g and Extended Data Fig. 4d). The responses of the GLI models were in good aggrement with the clinical results (Fig. 2g). P2, for instance, had a complete response (CR) after receiving four cycles of pembrolizumab combined with pemetrexed and cisplatin (PPEC). The corresponding GLI model demonstrated a high sensitivity (Ri = 0.734) to PPEC (Fig. 2f,g). The GLI model of P6, a SD patient with continuous tumor regression under pembrolizumab treatment combined with paclitaxel and carboplatin (PPAC), showed a moderate response (Ri = 0.542) to PPAC (Fig. 2f,g). Besides, a metastatic lesion with increased pleural fluid was discovered in P22 after two cycles of PPEC, and the P22-derived GLI model also showed no response (Ri = 0.169) to PPEC treatment (Fig. 2f,g).

Notably, the responses of LCO cultures without PBMCs were not consistent with the clinical results. For instance, the Ri of LCO cultures for the 3 SD patients were all less than 0.245 and categorized as NR (Fig. 2g). We further divided the 9 samples into 3 groups as double response (DR, Ri>0.245 for both the LCO and GLI cultures), single response (SR, Ri>0.245 for only the GLI culture) and no response (NR, Ri < 0.245 for both cultures). Interestingly, the DR, SR and NR groups corresponded to the clinical PR, SD and PD, respectively (Fig. 2g and Extended Data Fig. 4e). Pearson correlation coefficient (PCC) analysis demonstrated that the Ri of GLI correlated to the patient response better than that of LCO (Fig. 2h). In summary, the GLI model represented the clinical results of ICI therapy more accurately than LCO alone, suggesting the potential role of circulating T cells in anti-tumor immunity.

### Tumor-reactive T cells exist in the pre-treatment blood and reflect ICI-induced immune response

To further verify whether circulating T cells are tumor-reactive, we adopted the previously reported methods, activating the tumor-reactive T cells through suspension co-culture of patient-derived PBMCs with autologous tumor organoids.^20^ After 7-day co-culture, frequencies of CD8^+^ T cells expressing IFNγ and CD107a were upregulated in the R group (DR and SR samples), but not in the NR group (Fig. 3a,b). We then established GLI models using the post-activation T cells (CD8^+^ T cells sorted from suspension co-cultures, CD8^+^ T post) and autologous tumor organoids, and observed enhanced tumor cell death in the R group (Fig. 3c). Meanwhile, patient-derived PBMCs can neither be activated by nor kill the allogenic organoids (Fig. 3d-f), indicating that the organoid-induced activation of circulating T cells and the resulting cytotoxic effect are antigen-specific. In addition, PCC analysis indicated that Ri of the GLI model were significantly correlated with tumor reactivity of the circulating T cells (i.e., fold change of CD8^+^ T cells expressing either IFNγ or CD107a) but not total T cell frequencies (Fig. 3g-i). These results suggest the potential of circulating tumor-reactive T cells as an indicator of immunotherapy efficacy.^31^

**Fig. 3.**
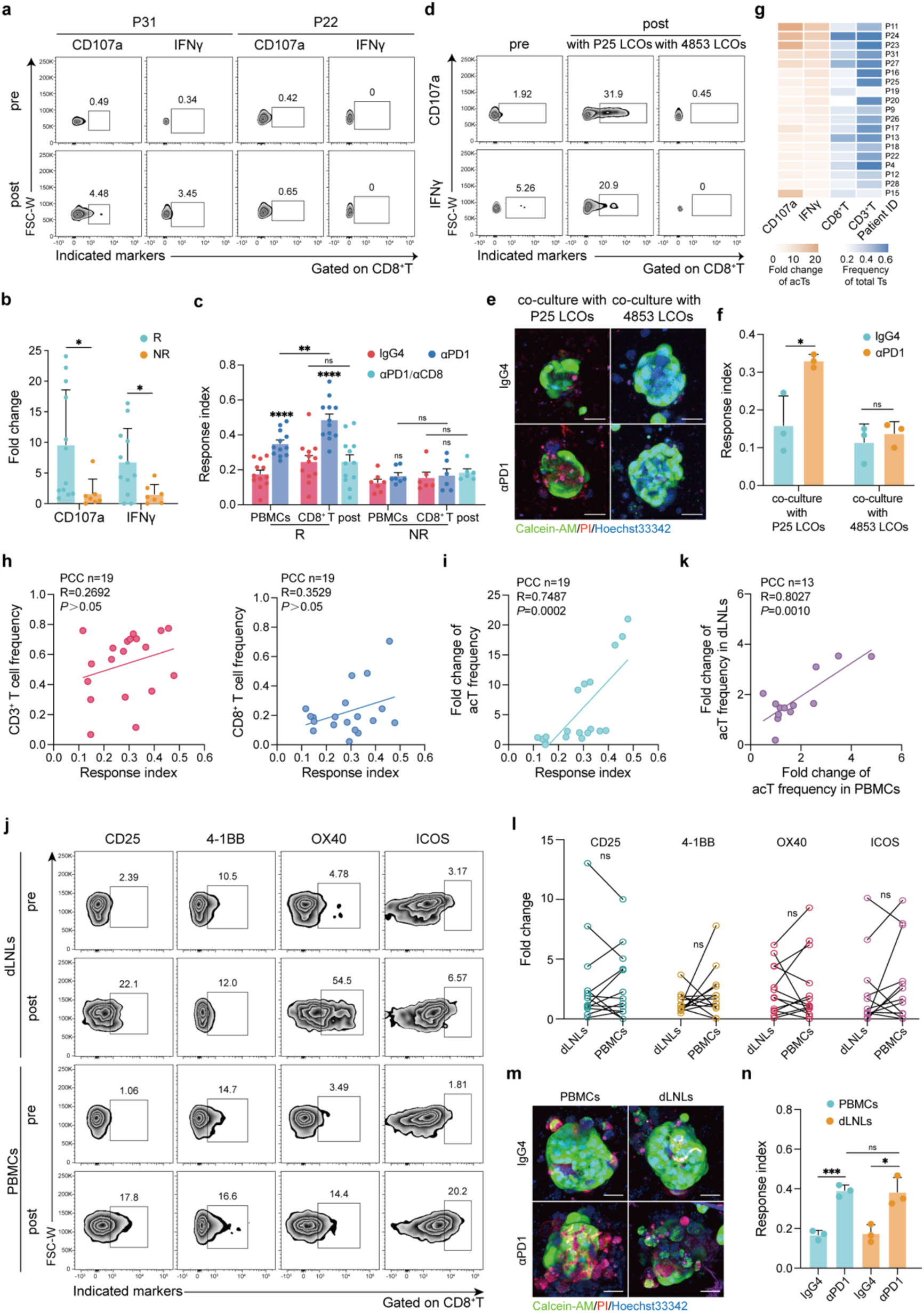
Tumor-reactive T cells exist in peripheral blood and correlate with ICI-induced immune response. **a,** FACS analysis of CD8^+^ T cells positive for CD107a or IFNγ in P31 and P22 PBMCs post 7-day suspension co-culturing with autologous LCOs. **b,** Bar plots comparing the fold change of CD107a or IFNγ positive CD8^+^ T cell frequencies following suspension co-culture (n=19). **c,** Bar plots comparing Ri of GLI models co-cultured with PBMCs or CD8^+^ T cells post suspension co-culture (CD8^+^ T post). **d,** FACS analysis of CD8^+^ T cells positive for CD107a or IFNγ in P25 PBMCs post 7-day suspension co-culture with P25 or 4853 LCOs. **e,** Fluorescence microscopy of P25 and 4853 LCOs in the GLI co-cultured with P25 PBMCs following 6-day IgG4 or αPD1 treatment. Scale bars: 50 μm. **f,** Bar plots comparing Ri of P25 and 4853 LCOs in the GLI co-cultures of (**e**) (n=3). **g,** Heatmap summary of CD3^+^ and CD8^+^ T cell frequencies in PBMCs from 19 patients and fold change of activated CD8^+^ T cells (acT, defined as CD8^+^ T cells positive for either CD107a or IFNγ) post 7-day suspension co-culture. **h,i,** PCC analysis between Ri of GLI models and CD3^+^ or CD8^+^ T cell frequencies in PBMCs (**h**) or fold change of acT frequency (**i**). **j,** FACS analysis of CD8^+^ T cells positive for activation markers (CD25, OX40, 4-1BB and ICOS) in PBMCs and dLNLs post 7-day suspension co-culturing with autologous LCOs. **k,** PCC analysis between fold change of acT frequency in dLNLs and in PBMCs. AcT defined as CD8^+^ T cells positive for one or more of the activation markers in (**j**). **l,** Fold change of CD8^+^ T cells positive for the indicated activation markers in PBMCs and dLNLs following suspension co-culturing with autologous LCOs (n=13). **m,** Fluorescence microscopy of P9 LCOs in the GLI models co-cultured with autologous PBMCs or dLNLs. Scale bars: 50 μm. **n,** Bar plots comparing Ri of P9 LCOs in GLI co-cultured with PBMCs or dLNLs following IgG4 or αPD1 treatment (n = 3). (\**P*<0.05, \*\**P*<0.01, \*\*\**P*<0.001, \*\*\*\**P*<0.0001, ns: no significance by unpaired student’s *t* tests)

The systemic anti-tumor immunity induced by neoadjuvant ICI has been reported in recent studies.^31^ Tumor-reactive T cells with new and pre-existing clonal types may be primed in the draining lymph nodes (dLNs) and then traffic through peripheral blood into the tumor tissue.^31–33^ Thus, we speculate that the tumor-reactive circulating T cells may be a reflection of the primed T cells in dLNs. To verify this speculation, we compared PBMCs and dLN-derived lymphocytes (dLNLs) on their potential of being activated by autologous tumor organoids using additional 13 patient samples from whom dLNs without metastasis were collected (Fig. 3j-l; Table 3). Post suspension co-culture with autologous tumor organoids, similar levels of T cell activation were observed in paired PBMCs and dLNLs as indicated by the levels of CD25, 4-1BB, OX40 and ICOS (Fig. 3j,l).^34^ We then compared the frequency of activated CD8^+^ T cells (acT) expressing any one of these markers. In 7 of the 13 samples, the fold changes of acT frequencies in PBMCs were higher than those in dLNLs, while in the another 6 samples, fold changes in dLNLs were higher (Fig. 3k). The PCC analysis indicated that the fold changes of acT frequencies in PBMCs were significantly correlated with those in dLNLs (R=0.8027, *P*<0.05) (Fig. 3k). We then built the GLI models by co-culturing LCOs with either PBMCs or dLNLs where similar levels of tumor cell death were observed (Fig. 3m,n). These results susggested that the tumor reactivity of circulating T cells may be a reflection of the systemic anti-tumor immunity. In aggrement with our data, clonal expansion of circulating T cells was reported in patients receiving ICI therapy and was demonstrated to predict tumor infiltration and clinical results.^35–37^

### Effector memory-like T cells in PBMCs infiltrated into LCOs and transform into effector-like tumor-reactive T cells under αPD1 treatment

To further identify the tumor-reactive T cells in peripheral blood and to dissect the dynamic responses of these cells to ICI, we conducted FascRNA-seq analysis, in which both the phenotypic data and single-cell transcriptome of the LCOs under various culture conditions were obtained (Fig. 4a). A total of 21,162 cells from 8 patient samples were sequenced and classified into 9 cell types through unsupervised clustering according to their gene expression patterns and 3,405 T cells were identified (Extended Data Fig. 5a,b). In order to characterize the 3,405 T cells precisely, we adopted a large T cell dataset^7^ of 47 lung cancer patients receiving αPD1 therapy to form a merged 154,254 T cell atlas with 9 subsets (Fig. 4b and Extended Data Fig. 7a). We first compared T cells in PBMCs (PBMC-Ts) from the R and NR groups that we defined above. Effector memory T cells (Tem) and proliferating T cells (Tprof) were the dominant subsets in PBMC-Ts from the R group, whereas PBMC-Ts from the NR group were mainly consisted of naive T cells (Tn) and central memory T cells (Tcm) (Fig. 4c,d). Notably, PBMC-Ts in the R group exhibited features of effector memory and neoantigen-specific circulating T cells reported previously^9,13^ with upregulated gene expressions related to stemness (*TCF7*) and migration (*ITGB2*, *ITGB7*) (Fig. 4e and Extended Data Fig. 6a,g).^13^ Gene ontology (GO) analysis of the differentially expressed genes (DEGs) revealed that the pathways related to T cell activation, chemokine production and signaling were enriched in PBMC-Ts of the R group (Extended Data Fig. 6b). Furthermore, we compared the expression of previously reported genesets of neoantigen-specific T cells between our PBMC-Ts and the reference data.^9^ Interestingly, PBMC-Ts in the R group had a positive correlation while those in the NR group had a negative correlation with the reference data (Extended Data Fig. 6h).^9^

**Fig. 4.**
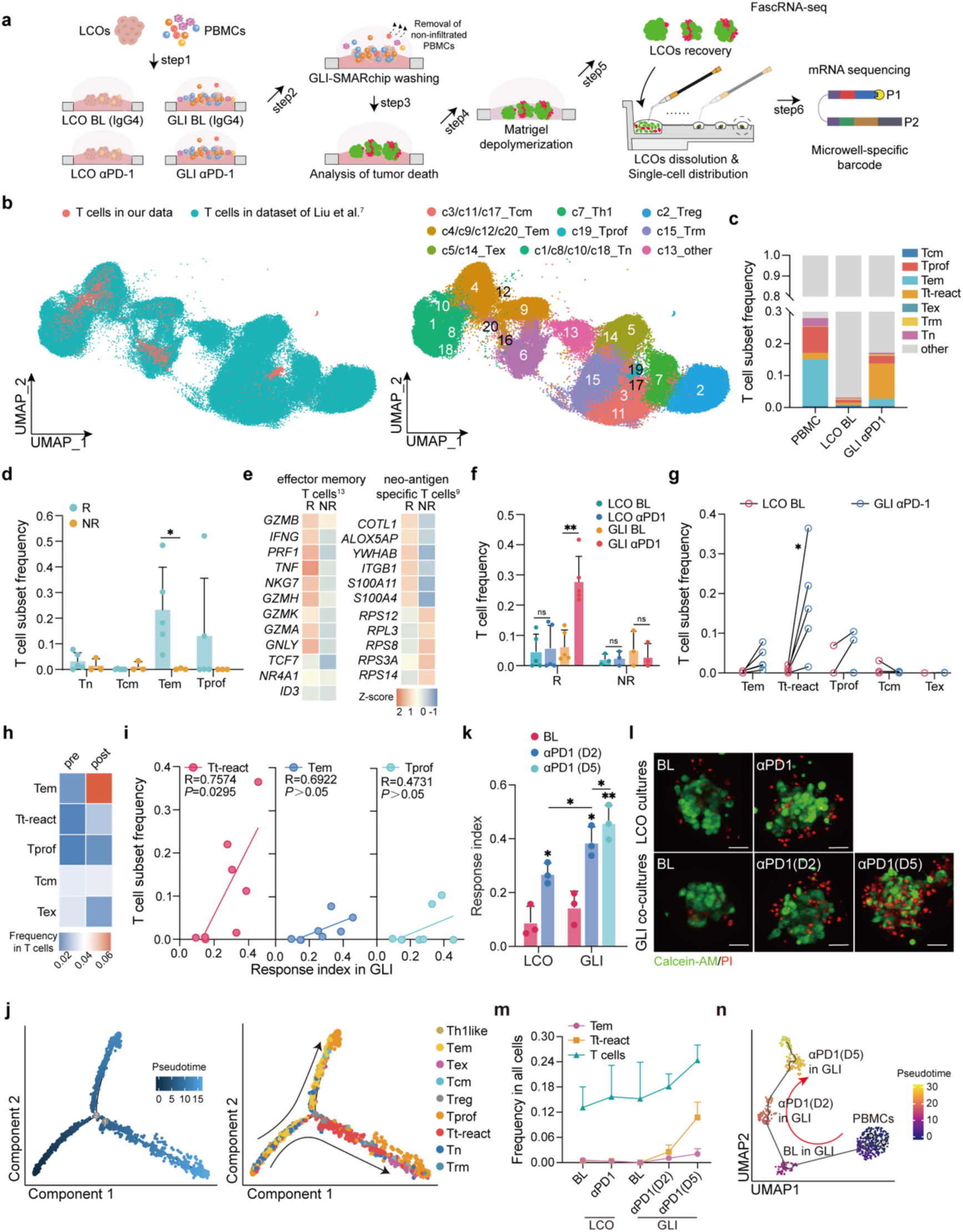
Tumor-reactive T cells in PBMCs are effector memory-like. **a,** Pipeline of the FascRNA-seq analysis of the LCO and GLI models. **b,** UMAP visualization of T cells from 8 lung cancer samples collected in our data combined with 47 lung cancer samples from the reference data^7^, labeled with respective sources (left) and cell types (right). **c,** Stack bar plots showing T cell subtype frequencies in PBMCs, LCO baseline (BL) and GLI αPD1 conditions. **d,** Bar plots comparing the T cell subtype frequencies in PBMCs from the R and NR groups. **e,** Expression levels of genes related to effector memory^13^ and neoantigen-specific^9^ features in PBMC-Ts from the R and NR groups. **f,** T cell frequencies in organoids harvested from the LCO and GLI co-cultures under all the experimental conditions. **g,** Dumbbell plots showing the T cell subtype frequencies in organoids cultured under the LCO BL and GLI αPD1 conditions for the R group, note the significant increase in Tem and Tt-react. **h,** Heatmap showing T cell subtype frequencies in the R group pre- and post-immunotherapy from the reference data^7^, note the increase in Tem and Tt-react. **i,** PCC analysis between Ri of GLI co-cultures and the frequencies of Tt-react, Tem and Tprof in the organoids recovered from the GLI co-cultures. **j,** Trajectory plots colored by pseudotime (left) and cell types (right), illustrating the inferred development trajectory. **k,** Bar plots showing Ri of P29 LCOs under different culture conditions (n=3). **l,** Fluorescence microscopy of P29 LCOs cultured in different conditions. Scale bars: 50 μm. **m,** Frequencies of total T cells, Tem and Tt-react in P29 LCOs cultured under different conditions (n=3). **n,** UMAP visualization plot displaying the inferred development trajectory by Monocle3. Arrow indicates the differentiation trajectory. (\**P*<0.05, \*\**P*<0.01, ns: no significance by unpaired student’s *t* tests)

Next, we explored the dynamic change of PBMC-Ts following the ecounter with tumor organoids in the GLI co-cultures. We first looked at whether PBMC-Ts can infiltrate into LCOs. As we expected, a significantly increased T cell frequency was observed in the GLI co-culture of the R group post αPD1 treatment while no change was seen in the LCO mono-cultures (Fig. 4f), suggesting the infiltration of PBMC-Ts into the LCOs, consistent with the observations in the MC38 GLI model. Furthermore, such phenomenon was not seen in the GLI co-culture of the NR organoids, indicating compromised interactions between LCOs and PBMCs (Fig. 4f). Specificly, Tem, Tprof and a specific subset of T cells (c6 in Fig. 4b) not identified in previous report^7^ increased significantly for all the R samples (Fig. 4g). Compared to Tem, the c6 T cells exhibited promoted features of effector and activation, while showing weaker exhaustion characteristics than exhausted T cells (Tex) (Extended Data Fig. 7a-d). Meanwhile, the similar trends were also seen in paired lung cancer samples before and after immunotherapy from the reference dataset (Fig. 4h).^7^ The PCC analysis revealed a strong correlation between the frequency of c6 T cells and Ri of the GLI models to αPD1 (Fig. 4i), suggesting that the c6 T cells were tumor-reactive and exerted the cytotoxic function. Therefore, we defined these cells as tumor-reactive T cells (Tt-react).

Interestingly, pseudotime analysis revealed a developmental trajectory of T cells initiated from Tn to Tem, then progressed to Tt-react and ultimately to Tex (Fig. 4j), in aggrement with the pseudotime analysis results of the reference data (Extended Data Fig. 7e,f).^7^ Genes related to cytotoxicity such as *NKG7, IFNG, CST7* and *GZMB* exhibited the highest expression levels in Tt-react, then decreased along the pseudotime trajectory, while T cell exhaustion markers such as *TIGIT, CTLA4, LAG3* and *CXCL13* were upregulated in Tex but not Tt-react (Extended Data Fig. 7g).^38^ Considering the low frequency of Tt-react in PBMCs, we speculate that Tem in PBMCs may infiltrate into LCOs and subsequently differentiate into the more activated and cytotoxic Tt-react to exert the tumor killing function.

To further confirm our speculation, we performed FascRNA-seq on P29 LCOs cultured in GLI at three time points (i.e., day 0, 2 and 5) post the αPD1 treatment. We found that the tumor cell death and the frequency of Tt-react continuously enhanced with time from day 0 (baseline, BL) to day 5 (D5) (Fig. 4k-m). Consistently, GO pathway enrichment demonstrated continuous upregulation of immune response (Extended Data Fig. 8a-d). In addition, the pseudotime analysis produced a development trajectory from PBMC-Ts to day 2 (D2) and then D5 in the GLI (Fig. 4n), we aligned with the real time series. The consistency between time series and pseudotime analysis demonstrates the reliability of the GLI model in tracking T cell dynamics. Overall, these data indicate that in response to ICI, Tem might infiltrate into tumor tissue, acquiring an effector-like state (Tt-react) and exerting the cytotoxic function.

### Circulating Tem with tumor-reactive potential are *CD9^+^CD44^+^GNLY^+^*

Since our data suggest the presence of Tem with tumor-reactive potential in peripheral blood, we then used bioinformatic methods to identify the gene expression signatures of these T cells. As shown in Fig. 5a, we first extracted the expression features of Tem before Tt-react along the pesudotime trajectory (Fig. 4j), obtaining 159 up-regulated (log2FC > 0.5) and 257 down-regulated (log2FC < −0.5) genes, mainly consisting of genes related to T cell function and activation (Extended Data Fig. 6c,d). Next, 120 T cell function-related genes were extracted from DEGs between the PBMC-Ts of the R and the NR groups, including 62 up-regulated (log2FC > 0.5) and 58 down-regulated (log2FC < −0.5) genes (Extended Data Fig. 6a). Then we took the intersection of these genes, leaving 48 up-regulated and 32 down-regulated genes (Fig. 5a).

**Fig. 5.**
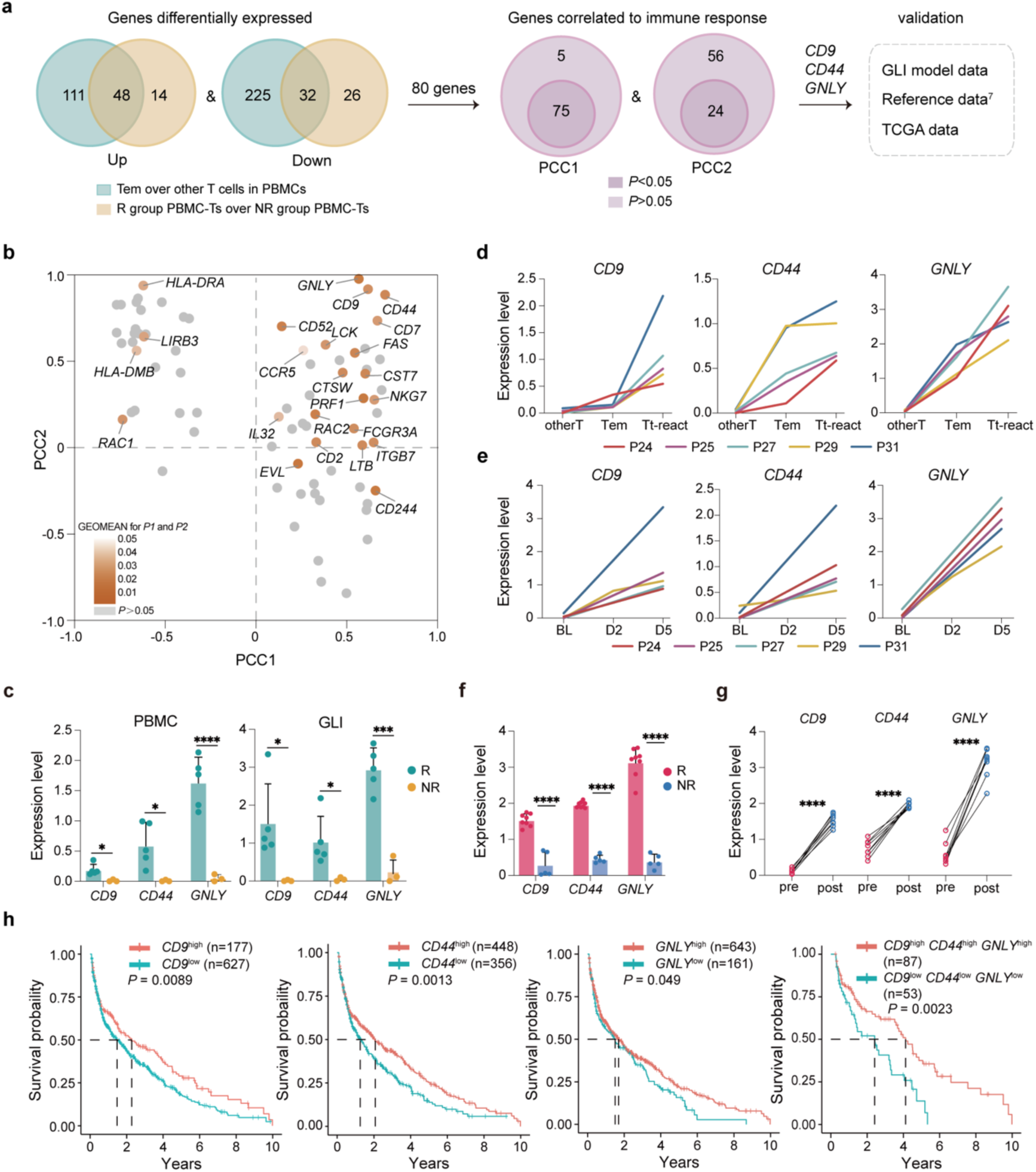
Signatures of Tem in pre-treatment blood with tumor-reactive potential. **a,** Schematic diagram illustrating the strategy for defining the signatures of Tem with tumor-reactive potential in blood. **b,** Dot plot showing the correlation between the expression level of the 80 genes identified from (**a**) and Ri of the GLI models under αPD1 treatment. PCC1 indicates the correlation between BL levels of gene expression in Tem and Ri, PCC2 indicates the correlation between Ri and gene fold change from Tem to Tt-react, note the correlations of *CD9*, *CD44* and *GNLY* are the most significant. **c,** Bar plots comparing the expression levels of *CD9, CD44* and *GNLY* in PBMC-Ts (left) and T cells harvested from GLI co-cultures (right) in the R and NR groups. **d,** Line charts showing the upregulation of *CD9, CD44* and *GNLY* in the R group, starting from other T cells in PBMCs to Tem and further to Tt-react in GLI models. **e,** Line charts showing the upregulation of *CD9, CD44* and *GNLY* in T cells for the R group, following αPD1 treatment. **f,** Expression levels of *CD9, CD44* and *GNLY* in tumor infiltrating T cells post-immunotherapy from the R and NR groups in reference data^7^. **g,** Expression levels of *CD9, CD44* and *GNLY* in tumor infiltrating T cells pre- and post-immunotherapy from the R group in reference data^7^. **h,** The Kaplan-Meier overall survival curves of LUAD and LUSD patients from TCGA database grouped by the expression levels of *GNLY*, *CD44 and CD9*. *P* value was calculated by multivariate Cox regression. (\**P*<0.05, \*\*\**P*<0.001, \*\*\*\**P*<0.0001 by unpaired student’s *t* tests)

Next, PCC analysis was performed to identify genes that were most significantly correlated with αPD1-induced tumor cell death. We first calculated the pearson correlation between the expression level of the 80 genes in Tem and Ri of the GLI co-culture under αPD1 treatment and 5 genes were eliminated owning to the nonsignificant correlation (Fig. 5a). Then, the fold changes of the 75 genes from Tem to Tt-react (FCT) were calculated and a second PCC analysis was performed between FCT and Ri (Fig. 5a,b). Based on the significance of the correlations and the dynamic changes in our time series measurements of the GLI models, *CD9, CD44 and GNLY* were identified as signatures of circulating T cells with tumor-reactive potential. CD44 is best known as a critical regulator of immune response, especially potentiating T cell activation and maintenance of memory cells.^39^ GNLY is mainly produced by activated CD8^+^ T cells and associated with cytotoxicity.^40^ CD9, a co-stimulatory signal of CD28, also plays an important role in T cell activation and migration.^41–43^

We validated the expression levels of the 3 genes in the R and the NR groups. In both PBMC-Ts and T cells from the GLI co-cultures, the expression levels of *CD9, CD44,* and *GNLY* were significantly higher in the R group than those in the NR group (Fig. 5c). Among the 5 samples responsive to αPD1, we similarly observed higher expression levels of *CD9, CD44* and *GNLY* in Tem compared to other T cells from PBMCs, and a further increase upon transition to Tt-react (Fig. 5d). Further, their expression levels following αPD1 treatment were consistently higher than BL in all the 5 samples (Fig. 5e), suggesting upregulation of the 3 genes were associated with responsiveness to αPD1 treatment. Additionally, in the reference data,^7^ compared to patients from the NR group, patients from the R group exhibited higher expression levels of all the 3 genes, which were also significantly elevated compared to pre-treatment levels (Fig. 5f,g). Moreover, analysis of the independent lung adenocarcinoma (LUAD) and lung squamouscarcinoma (LUSD) cohorts from The Cancer Genome Atlas (TCGA) indicated that elevated expression of *GNLY*, *CD9* and *CD44* were associated with significantly better overall survival (OS) (Fig. 5h). In conclusion, the presence of *GNLY*^+^*CD9*^+^*CD44*^+^ Tem cells in PBMCs may associate with tumor-reactive potential and could serve as predictive signatures for responsiveness to the αPD1 treatment.

### The LCO and GLI co-culture hint the local and systemic anti-tumor immunity

In the local tumor micro- and the systemic macro-immune environment work together to define the response to immunotherapies.^33^ The LCOs and GLI co-culture models illustrated two patterns of αPD1-induced anti-tumor immunity in lung cancer: i) both the tumor infiltrated and circulating T lymphocytes can be activatied to kill the tumor cells (i.e., both LCO and GLI responded to αPD1, defined as DR), and ii) only the circulating T cells can be activated (i.e., only GLI responded to αPD1, defined as SR). To clarify whether these two patterns correspond to different local and systemic immune status, we further analyzed the single-cell transcriptome data of these two groups. Although the frequency of T cells were similar in the PBMCs for both groups (Fig. 6a), the DR group has more Tprof (Fig. 6b) with elevated expressions of T cell stemness and proliferation genes, including *TCF7* and *MKI67* (Fig. 6c and Extended Data Fig. 6e). Whereas the SR group was dominant with Tem (Fig. 6b), along with upregulated expressions of genes related to T cell cytotoxicity and activation such as *IFNG*, *GZMB* and *CD69* (Fig. 6c and Extended Data Fig. 6e). The DEGs between PBMC-Ts in these two groups were enriched to inflammatory signaling pathways, including the JAK-STAT and NF-κB pathways (Extended Data Fig. 6f), suggesting a more inflammatory status for immune cells in the SR group.

**Fig. 6.**
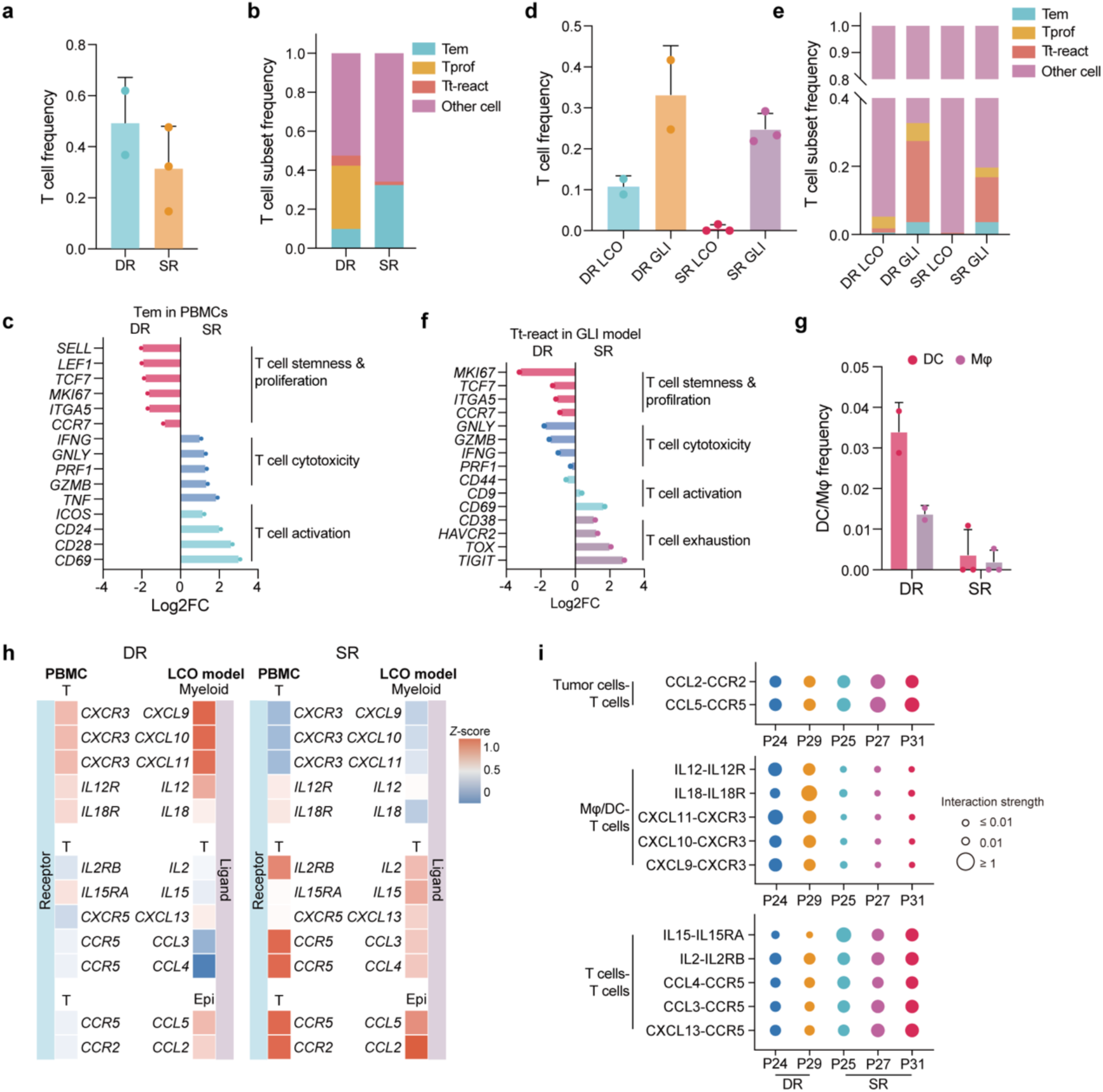
T cells in PBMCs infiltrate into LCOs through different chemotactic paths in DR and SR groups. **a,** Bar plots showing T cell frequencies in PBMCs for the DR and SR groups. Each dot represents a patient sample. **b,** Stack bar plots showing frequencies of Tem, Tt-react, Tprof in PBMCs from the DR and SR groups. **c,** Fold change of indicated genes in Tem of PBMCs between the SR group and the DR group. **d,** Bar plots showing T cell frequencies in organoids cultured under different conditions in the DR and SR groups. **e,** Stack bar plots showing frequencies of Tem, Tprof and Tt-react in organoids cultured under different conditions for the DR and SR groups. **f,** Fold change of indicated genes in Tt-react of GLI models between the SR group and the DR group. Note the elevated expression of exhaustion marker genes in the SR group. **g,** Bar plots showing frequencies of DC and Mφ in LCO BL for the DR and SR groups. **h,** Expression levels of selected chemokine and cytokine receptors in PBMC-Ts and the corresponding ligands in T cells (T), myeloid cells (including Mφ and DC) and epithelial cells (Epi) of the autologous LCOs. Note chemotactic interaction between PBMC-T and LCO-myeloid cells is most significant for the DR group, while the interaction between PBMC-T and LCO-T or LCO-Epi are the most obvious for the SR group. **i,** Dot plots showing the strength of intercellular communication of indicated ligand-receptor pairs in the DR (P24/29) and SR (P25/27/31) groups.

In the GLI co-cultures, the T cell frequencies, especially the Tt-react, in the LCOs increased significantly in both the DR and SR groups post αPD1 treatment (Fig. 6d,e), suggesting an efficient T cell infiltration. The gene expression analysis indicated Tt-react from the DR group exhibited intensified stemness, proliferation and cytotoxicity features compared to the SR group (Fig. 6f), yet showing less exhaustion features (Fig. 6f). These results suggested that PBMC-Ts of the SR group underwent exhaustion more rapidly than those of the DR group, probably owning to their more inflammatory state. Accordingly, continuous replenishment of new tumor-reactive T cells from the blood is needed to maintain a long lasting anti-tumor effect, consistent with the phenomenon that LCOs from the SR group had less T cell infiltration and did not respond to ICI when cultured alone.

Besides the different states of PBMC-Ts, the local tumor immune microenvironments (TIMEs) also exhibited distinct features, thus leading to diverse pathways for T cell recruitment in the two groups. More myeloid cells, including macrophages (Mφ) and dendritic cells (DC), were observed in LCOs of the DR group (Fig. 6g). We classified Mφ and DC according to previously reported features of their subsets and identified Mφ-CCL2, mDC and cDC2-FCGR2B subtypes (Extended Data Fig. 9a-d), which were reported to promote lymphocytes infiltration.^13^ These myeloid cells exhibited elevated levels of chemokine and cytokine expression, including *CXCL9*, *CXCL10, IL12* and *IL18*, and the corresponding receptors were upregulated in PBMC-Ts of the DR group compared to the SR group (Fig. 6h). Cell-cell interaction analysis indicated T cells in the SR group were mainly recruited through the *CCL5-CCR5* and *CCL3-CCR5* axis of T-T cells and epithelial-T cells, while typical *CXCL9/10/11* related interactions of myeloid-T cells were the major paths for T cell recruitment in the DR group (Fig. 6i and Extended Data Fig. 9e,f).^44^ In summary, these data demonstrated the responses of LCO mono-culture and GLI co-culture models to ICI are correlated to the characteristics of the local TMEs and the systemic immune environments.

## Discussion

In recent years, there has been growing interests in understanding the systemic relationship between blood, dLNs and tumors, as the alternative sources of functional effector T cells are necessary for sustained anti-tumor immunity under ICI treatment.^31^ These exogenous immune supplies rely on the integrity of peripheral immune function, communication, and trafficking. ICI increases the interactions such as CD28 co-stimulation between traditional DCs and Tn in the dLNs,^45,46^ leading to rapid expansion of T cell clones specific to tumor-associated antigens and their circulation into the blood.^33,36,37^ These expanding peripheral T cells eventually infiltrate into tumor tissues, expressing markers related to antigen-specific activation and exert cytotoxic function.^36,37^ Researches have emphasized the presence of tumor-reactive T cells in the blood,^9,20^ which were activated and underwent clonal expansion during ICI treatment,^7,31^ targeting various tumor antigens.^9,47^ In this study, we co-cultured immune cells obtained from both PBMCs and dLNs with LCOs using the GLI model in parallel. The similarities in tumor killing and T cell activation between the two groups suggested an interconnection between dLN and peripheral blood in lung cancer patients. Therefore, we believe circulating T cells in peripheral blood can be an indicator of the systemic anti-tumor immunity as their tumor reactivity and status are highly correlated to tumor regression under ICI treatment.^48^

The GLI model not only preserved naïve immune cells within PDOs but also incorporated circulating PBMCs, thus providing a more comprehensive *ex vivo* tumor model containing at least part of the systemic immune components compared to current strategies.^18,26^ The GLI co-culture enabled efficient migration of immune cells towards LCOs as a result of chemokine ligand-receptor interactions, thus mimicing the *in vivo* immune cell infiltration (into the tumor tissue) as well as the subsequent TCR-dependent CD8^+^ T cells activation and fate transformation. In addition, the low cost of GLI models and their convenience in operation and analysis enabled fully dissection of the dynamics of anti-tumor immunity under various conditions, such as the ICI treatment in the current study.

Cell therapy has emerged as a promising approach in cancer treatment, as the first TILs treatment is hitting the clinic.^49^ Be that as it may, obtaining tumor-specific and tumor-reactive T cells as well as their TCRs remains challenging.^50,51^ In this one as well as our previous study, we have demonstrated the feasibility of identifying tumor-reactive T cells through a combined phenotypic-transcriptome analysis of LCO cultures.^22^ In addition, we identified *GNLY*, *CD44* and *CD9*, a co-stimulatory signal of *CD28*, as potential signatures of tumor-reactive T cells in peripheral blood.^41,52^ In the future, these identified gene signatures can be used to diagnose or even isolate tumor-reactive T cells from blood as cell therapy products.

Great efforts have been made to identify tumor-specific T cell clonotypes, including a peripheral blood lymphocytes (PBL) derived neoantigen-specific T cell discovery strategy using tetramers of neoantigens,^9^ as well as the PDO-based co-culture system for the expansion of tumor-specific T cells from PBMCs.^20,53^ While these approaches ensure the specific recognition of tumor antigens, the states of T cells were overlooked, i.e., states of T cells corresponding to efficient tumor infiltration and sustained cytotoxicity need to be clarified in order to develop an effective cell therapy strategy for solid tumors. Interestingly, our scRNA-seq data of the GLI co-culture suggested stemness (high level of *TCF7, MKI67*) of T cells are correlated with more efficient activation and tumor killing while elevated inflammation signals indicate susceptibility to exhaustion and compromised tumor tissue infiltration, suggesting a correlation between inflammation and immune suppression which worth further exploration.

## Methods

### Human material

All human lung cancer tissues, peripheral blood, and draining lymph nodes were obtained from patients with a confirmed pathological diagnosis of lung cancer. These patients underwent surgical resection or biopsies at Peking University People’s Hospital. All experiments utilizing human material were approved by the ethnical review boards of Peking University People’s Hospital and Tsinghua University. Written informed consent for research was obtained from donors prior to tissue acquisition. Detailed sample information is available in Table 4.

### GLI-SMARchip fabrication

The fabrication process of the GLI-SMAR chip is illustrated in Extended Data Fig. 1b. Briefly, silica-polyethylene terephthalate (silica-PET) double-side tape was purchased from Adhesive Research (York, USA) and laser-cutted into microarray with a diameter of 1.2 mm and inter-well spacing of 2.25 mm. The superhydrophobic layer is fabricated according to our previous report.^23^ One side of the protective film was removed and the slide with the superhydrophobic layer was pressed against the microwell array tape for 10 minutes. Upon separation, a 40-μm-thick superhydrophobic layer was grafted onto the surface of one side of the microwell array tape, providing physical isolation. A thin layer of Dow Corning 3140 was spin-coated on a 4′′ PMMA (poly (methyl methacrylate)) wafer at 7000 rpm for 30 seconds and then transferred to the opposite side of the tape by contact printing after removal of protective film. Subsequently, the tape with superhydrophobic layer was transferred to the glass bottom of a confocal dish, forming a GLI-SMARchip.

### Construction and optimization of GLI co-culture model

The GLI co-culture pipeline is illustrated in Extended Data Fig. 1c. Firstly, Matrigel was diluted to 80 % concentration using LCO culture medium (LCOM, listed in Table 4). Then, 0.6 μL Matrigel containing 5-10 LCOs were dispensed into each microwell of the pre-cooled GLI-SMARchip with a pipette at 4°C. Next, the GLI-SMARchip was inverted at 4°C for 10 minutes and subsequently transferred to room temperature (RT) while still inverted. After 5 minutes, once the Matrigel had solidified, 2 μL of T cell expansion medium (TCEM, STEMCELL) with or without PBMCs were added to the microwells for droplet culturing.

The optimization of the GLI co-culture model focused on two key parameters: inverting time at 4°C and Matrigel concentration. The optimized parameters were determined as follows: We quantified the percentage of LCOs located between 180 μm to 220 μm in the Z-axis direction relative to the total number of LCOs in the microwells of the GLI-SMARchip at different Matrigel concentrations (50%, 60%, 70%, 80%, and 90%) and inverting times (5 minutes, 10 minutes, and 15 minutes) at 4°C. A 10-minutes inverting time and 80% Matrigel concentration resulted in the highest number of LCOs at the indicated Z-position (Extended Data Fig. 2b).

### Syngeneic murine models

MC38 and LLC cells (1×10^6^ cells per mouse in 100 μL PBS) were injected into 8-week-old female C57BL/6 mice, and tumors were collected one to two weeks later upon reaching a size of 1000 mm^3^. For the *in vivo* immunotherapy studies, mice were randomized and injected with 10 mg/kg of either isotype control IgG (BioXCell) or αPD1 (BioXCell) every 3 days for 7 doses. Tumor volumes were measured using vernier calipers throughout the treatment period.

### Generation, culture and drug treatment of MDOs and LCOs

Fresh tumor specimens of mouse or human patients were received in sample collection media (DMEM + 10 % FBS+ 1 % penicillin-streptomycin for mouse and Advanced DMEM/F12 + 1 % Penicillin-streptomycin for patients) on ice and minced in a 10-cm dish using sterile forceps and scalpel. The minced tumor was then pelleted and resuspended in fresh media before being strained through 100 μm f and 40 μm filters to generate S1 (>100 μm), S2 (40-100 μm), and S3 (<40 μm) fractions. S2 fractions were collected and cultured in ultralow-attachment plates and formed MDOs or LCOs after overnight suspension. The following day LCOs or MDOs in suspension were centrifuged for 5 minutes at 2000 rpm in 4°C and resuspended in the cold growth factor-reduced Matrigel. Subsequently, 60 μL drops of the Matrigel cell cluster suspension were inoculated into ultralow-attachment 96-well tissue culture plates with a flat bottom and allowed to solidify at 37°C for 20 minutes. Once the Matrigel became stable, cells were overlaid with respective medium: DMEM + 10 % FBS+ 1 % penicillin-streptomycin for MDO and LCOM for LCOs. For experiments on the GLI-SMARchip, 0.6 μL Matrigel containing 5-10 organoids was loaded into each microwell with a pipette. Each Matrigel droplet in the microwell was then overlaid with 2 μL TCEM with or without indicated therapeutic monoclonal antibody using the spot-cover method. MDOs were treated with isotype control IgG (10 μg/mL), αPD1 (10 μg/mL) or αPD1 (10 μg/mL) + αCD8 (10 μg/mL). LCOs were treated with isotype control IgG4 (10 μg/mL), αPD1 (nivolumab, 10 μg/mL), combination treatment (10 μg/mL nivolumab + 10 μM pemetrexed/paclitaxel + 10 μM cisplatin/carboplatin) or reversion treatment (10 μg/mL nivolumab + 10 μg/mL αCD8).

### Isolation of PBMCs from blood and lymphocytes from dLN

Venipuncture is used for obtaining blood samples from humans, while retro-orbital sinus bleeding is used for collecting blood from mice. PBMCs were isolated from peripheral blood by Lymphoprep^TM^ density gradient separation and cultured in TCEM. Lymphocytes from dLN were isolated by FACS. Single cell suspensions were centrifuged, resuspended in 1 mL of FACS wash buffer (FWB; PBS + 2% FBS + 1 mM EDTA), and stained with antibodies including Human Fc Block (BD), CD45-Pacific Blue (BioLegend), and near-infrared viability dye (Invitrogen). Following a 30-minute incubation at 4°C in the dark, cells were incubated on ice for 20 minutes, washed with 1 mL FWB, and then resuspended in 5 mL FWB. CD45^+^ immune cells were sorted using a FACS Aria II or III instrument (BD) and immediately transferred to ice. Lymphocytes were cultured in TCEM.

### Flow cytometry analysis

Tumors from MC38 and LLC syngeneic mouse models were procured as described above and single cells were prepared by dissociation with 300 units/mL collagenase IV (Sigma-Aldrich) at 37°C for 30 minutes, followed by three 5-minutes washes with FWB. For MDOs or LCOs cultured on the GLI-SMARchip, PBMCs that did not infiltrate into the MDOs were removed by washing with PBS for two times. Each microwell was then overlaid with 2 μL of organoid digestion mix including Organoid Harvesting Solution (R&D Systems) with 10 mg/mL *B. Licheniformis* protease (Sigma-Aldrich) and incubated on ice for 1 hour. The LCOs in the microwells were then harvested using a pipette and digested into single-cell suspensions using TrypLE (Gibco) followed by three 5-minutes washes with FWB.

For mouse samples, FcR were blocked by incubation with the anti-mouse CD16/CD32 blocking Ab (BioLegend) for 15 minutes at 4°C. Cell-surface staining was performed using the following antibodies: CD45-PE Cy5 (BioLegend), CD3-Alexa Fluor 700 (BioLegend), CD4-FITC (BioLegend), CD8-BV421 (BioLegend), and near-IR viability dye (Invitrogen) for 30 minutes at 4 °C. For human samples, FcR were blocked prior to surface antibody staining using Human FcR Blocking Reagent (BD) for 10 minutes at RT. Cell-surface staining was performed using the following antibodies: CD45-Pacific Blue (BD), CD3-BV510 (BD), CD4-FITC (BD), CD8-PerCP-Cy5.5 (BD), and near-IR viability dye (Invitrogen) for 30 minutes at 4 °C. After cell-surface staining, cells were fixed and stained for intracellular IFNg-APC (BioLegend for mouse; BD for patients) using the Cytofix/Cytoperm kit (BD), according to manufacturer’s instructions. CD107a-PE antibodies (BioLegend for mouse; BD for patients) were added before cell-surface staining and Protein Transport Inhibitor Cocktail (1X, eBioscience) was added after 1 h and continued for an additional 4 h. Cells were washed and resuspended in FWB and analyzed using a FACS Aria II or III instrument (BD). Data were analyzed using FlowJo software version 10.8.1.

### Quantitative real-time PCR

Analysis of expression levels of *IFNg*, *Prf1* and *Gzmb* by quantitative reverse transcription polymerase chain reaction (qRT-PCR) was performed using CD3^+^T cells sorted from mouse TT and GLI models. Tissue samples were snap-frozen in liquid nitrogen and processed to extract RNA, which was stored at −80°C after extraction. Primers were designed for *IFNg* (Fwd: 5′-AAGACAATCAGGCCATCAGCA-3′, Rev: 5′-TGGTGGACCACTCGGATGA-3′), *Prf1* (Fwd: 5′-CTCCACGCATGATCTGCTCT-3′, Rev: 5′-CACCGGGCTCTGCTCATTAT-3′), *Gzmb* (Fwd: 5′-GCGCAATGTCAATGTGAAGC-3′, Rev: 5′-GCACGTTTGGTCTTTGGGTC-3′) and *Gapdh* (Fwd:5′-CATCACTGCCACCCAGAAGACTG-3′, Rev: 5′-ATGCCAGTGAGCTTCCCGTTCAG-3′). *Gapdh* was used as reference gene to normalize the expression of target genes. Total RNA was extracted via the QIAGEN RNeasy Mini Kit. qRT-PCR was performed on a StepOnePlus instrument (Applied Biosystems) with an annealing/extension temperature of 60 C for 35 cycles. Paired Student’s *t-test* was employed to calculate P values for statistical analysis of gene expression levels.

### Immunofluorescence and quantitative evaluation of drug sensitivity

To perform PBMC infiltration tracking, LCOs were in some cases stained with 1 mM of CellTrace CFSE (Invitrogen) and T cells with 100 nM CellTrace FarRed (Invitrogen) in PBS for 20 min at 37°C followed by blocking with FBS and washing in PBS. LCOs in the GLI model were imaged daily using a 40X silicone oil objective on a Nikon A1HD25 confocal microscope.

For viability assessment, LCOs and MDOs were washed with PBS and stained using the Calcein-AM/PI Double Stain Kit (Invitrogen) along with Hoechst 33342 (10 μg/mL, Invitrogen). Following incubation with the dyes for 30 minutes at 37°C in the dark, images were captured using a 10X objective on a Nikon A1HD25 confocal microscope. Image capture and analysis were performed using the NIS-Elements AR software package. Initially, all tumor cells were identified based on Hoechst 33342 staining and cell size (12-13 μm). Cells stained with both Hoechst 33342 and Calcein-AM were classified as viable, while cells stained with Hoechst 33342 and PI were considered dead. The response index was calculated by determining the number of dead tumor cells relative to total tumor cells.

### Single-cell preparation and transcriptome library construction

FascRNA-seq was conducted following previously described methods.^22^ Briefly, a library of 384 capture oligos with unique barcodes was prepared. Single cells were loaded into the reaction chip using the single-cell distribution instrument (SCDI). A transfer coverslip containing magnetic beads and barcoded capture oligos was then aligned and placed onto the reaction chip. The assembled MoSMAR-chip was washed with 5X RT buffer, and single cells were lysed *in situ* with lysis buffer for 15 minutes. mRNA captured by the magnetic beads was subsequently washed and resuspended in 5X RT buffer for downstream processing.

After loading with dNTPs, Template Switch Oligo (TSO), Reverse Transcriptase, and RNase inhibitor, the mixture was incubated at room temperature for 30 minutes followed by 42°C for 90 minutes to generate cDNA. Subsequently, PCR was conducted by adding 2× HotStart Readymix (Kapa Biosystems), 5′-end biotin-modified P7 primer (10 μM), and TSO primer (10 μM). The PCR products were purified using 0.6× AMPure XP beads (Beckman Coulter) and eluted in 50 μl ddH2O. The PCR products were fragmented using the Covaris M220 system (Covaris) to generate 3′-end cDNA fragments. Subsequently, end repair, 5′ phosphorylation, dA-tailing, adaptor ligation, size selection, and PCR enrichment were performed sequentially using the NEBNext Ultra™ II DNA Library Prep Kit for Illumina (NEB). Finally, PCR products were purified and the corresponding scRNA-seq library was applied quantity control using the Agilent Bioanalyzer 4200. The libraries were subjected to paired-end sequencing (150 nt each) on the HiSeq-PE150 instrument (Illumina). Two paired-end reads were obtained: Read 1 containing a sequence typically mapping to the 3′ end of an mRNA transcript, and Read 2 containing a wellcode (16 bases) identifying a specific microwell along with a unique molecular identifier (UMI) (10 bases).

### Single-cell RNA-seq analysis of tumor organoid models

Dataset we referred was obtained from Gene Expression Omnibus databases (GSE179994).^7^ The bioinformatics analysis pipeline utilized for this study is available on Github. Firstly, Read 1 Fastq files were aligned to the reference genome using STAR v2.4.0a with default parameters. Read 2 Fastq files were processed to extract wellcodes and unique molecular identifiers (UMIs) based on the wellcode-UMI-Poly T pattern (16-nt wellcode, 10-nt UMI, 16-nt poly T).

In downstream data analysis, we applied quality control on scRNA-seq data with >300 detected transcripts and <20% mitochondrial reads to retain. Filtered single-cell transcriptomic data was analyzed using Seurat version 4.2.0.^54^ The data was normalized and variable genes were identified using the NormalizeData and FindVariableFeatures functions (selection method = “vst”, features = 1500). Harmony algorithm was employed for batch effect correction based on patient samples, and the integrated dataset was scaled and used for principal component analysis (PCA).^55^ UMAP dimensional reduction and cluster identification were performed using the first 10 principal components. Differential gene expression analysis was conducted using FindAllMarkers function. Cluster-specific pathways were determined using KEGG and GO enrichment analysis via clusterProfiler version 4.4.4.^56^ Cell type annotations were firstly performed by SingleR (version 1.10.0) with BlueprintEncode and HumanPrimaryCellAtlas as reference datasets.^57^ Integrated cell types were then refined by DEGs of cluster-specific expression features, enabling comprehensive cell type identification across the dataset.

The R package Monocle 2 (version 2.30.0) and Monocle 3 (version 1.3.1) was used to explore pseudotime trajectories among T cells from scRNA-seq data.^58,59^ For T cells from our dataset, we applied Monocle 2 to derive pseudotime trajectory in the form of DDRTree with default parameters. For dataset of merged total T cells, we adopted Monocle 3 due to the limited data processing throughput and calculating trajectories in UMAP visualization also with default parameters. The normalized expression level of concerned genes was further plotted along the pseudotime trajectory.

Cell communication analysis was conducted with CellCall (version 0.0.0.9000) and CellChat (version 1.6.1) R packages.^60,61^ We identified overexpressed gene combinations and interactions using identifyOverExpressedGenes and identifyOverExpressedInteractions, followed by pathway analysis using computeCommunProb and netAnalysis_computeCentrality in Cellchat. In Cellcall analysis, we applied CreateObject_fromSeurat to create data object and use TransCommuProfile to infer intercellular communication by integrating ligand/receptor expression and downstream TF activities. Results from above were aggregated and subjected to further statistical analysis.

### TCGA data analysis

To investigate the correlation between selected genes and patient survival, data from TCGA were utilized. Gene expression data and corresponding clinical information were retrieved from LUSC and TLUAD projects. For each selected gene within a given TCGA ID, the highest expression level was retained for subsequent Kaplan-Meier survival analysis. Concurrently, clinical data were processed to obtain survival time data for each ID. When performing statistical analysis, the surv_cutpoint function (from R package “survminer”) was used to provide cutpoints that correspond to the most significant relation with the outcome (here, survival), and “high expression” and “low expression” groups were categorized based on the given cutpoints. Curves were fitted using survfit, and the significance of survival differences for each selected gene was tested using survdiff to calculate P values.

### Statistical Analysis

GraphPad Prism 10 was used to plot and statistically analyze the data. The differences between two groups were analyzed by unpaired, two-tailed student’s *t* tests. The significances were denoted as * and ns, (specifically, \**P* < 0.05, \*\**P* < 0.01, \*\*\**P* < 0.001, \*\*\*\**P*<0.0001, *P* value lower than 0.05 is considered significantly different). Values in column bar plots were illustrated as mean ± standard deviation (SD). Transcriptome related data process procedure was presented in experimental sections above. Flow cytometry data were analyzed with FlowJo software. Image J was used for image signal processing and quantitative statistics.

---

**Extended Data Fig. 1.**
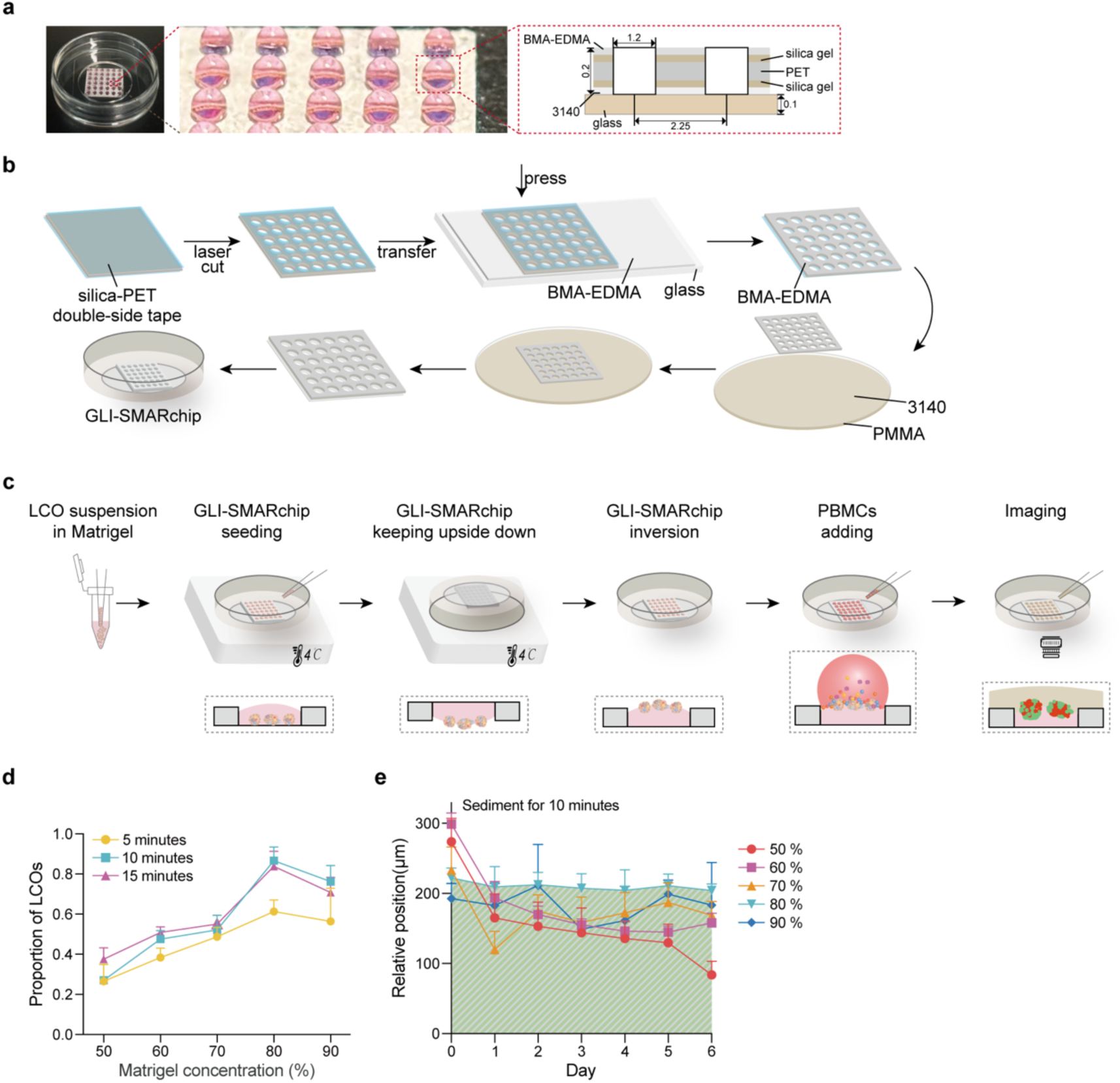
The GLI-SMARchip and optimization of the GLI co-culturing procedure. **a,** Image of the GLI-SMARchip displaying an array of 36 microwells and the cross-section of the chip (unit: mm). **b,** Micrografting procedure for the GLI-SMARchip fabrication. **c,** Operation procedure of the GLI co-culture model. **d,** Optimization of sediment time at 4°C and Matrigel concentration in the GLI co-culture model (n=5). **e,** Relative positions of LCOs in a well under different Matrigel concentrations over time (sediment for 10 minutes) (n=3).

**Extended Data Fig. 2.**
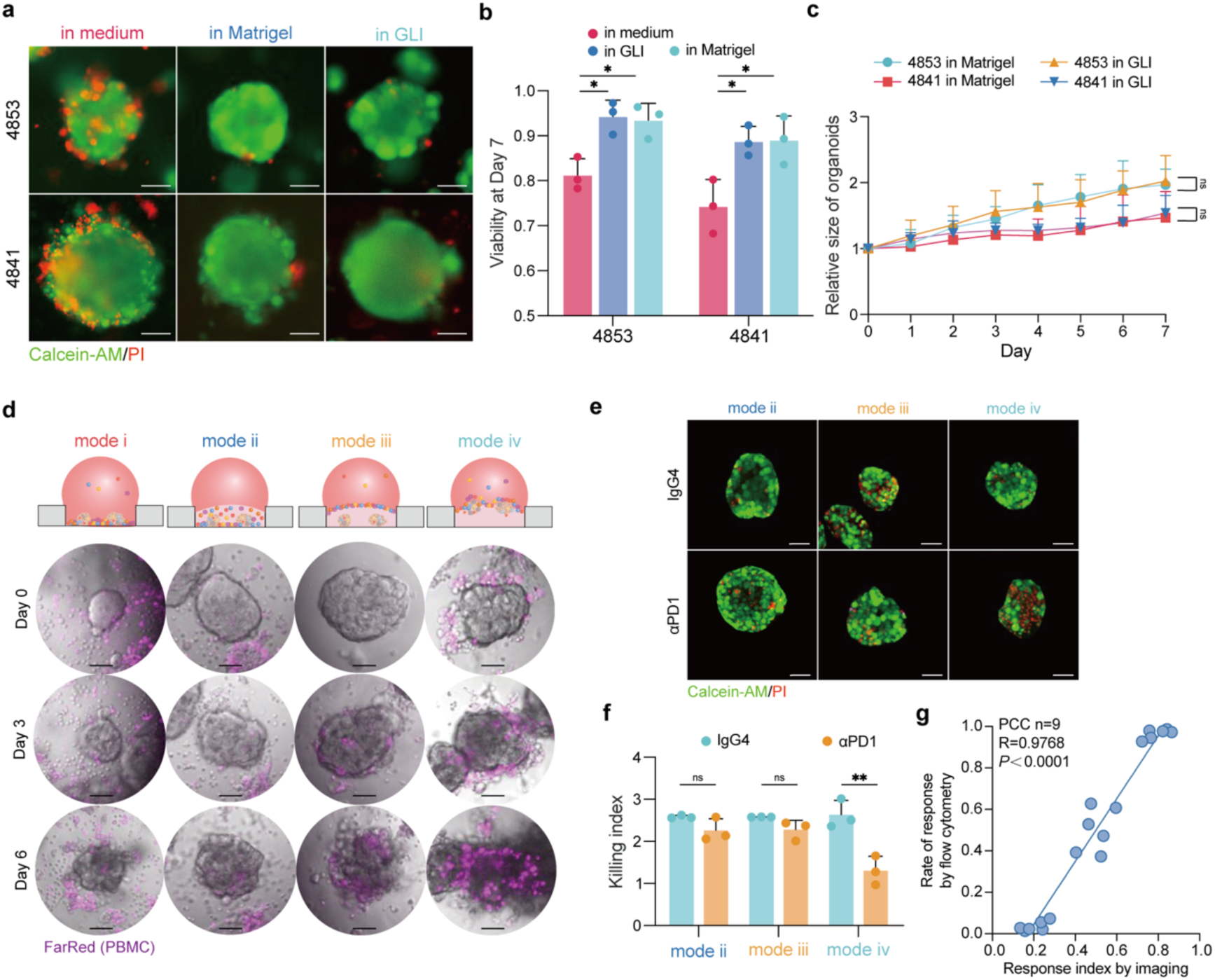
Characteristics of GLI co-culture model. **a,** Fluorescence microscopy of LCO 4853 and 4841 at day 7 showing the viability of LCOs cultured in medium, in Matrigel or in GLI. Scale bars: 100 μm. **b,** Quantification of viabilities for the three conditions in (A) (n=3). **c,** Comparison of growth rates for LCO 4853 and 4841 in Matrigel or in GLI over time (n=10). **d,** Representative 2D images showing PBMCs migrating and infiltrating into LCOs under different co-culture modes. Scale bars: 50 μm. **e,** Fluorescence microscopy showing viabilities of LCOs on day 6 under different co-culture modes. Scale bars: 50 μm. **f,** Killing index of LCOs following IgG or αPD1 treatment under different co-culture modes (n = 3). **g,** PCC analysis between imaging-based Ri quantification and flow cytometry-based quantification. (\**P*<0.05, \*\**P*<0.01, ns: no significance by unpaired student’s *t* tests)

**Extended Data Fig. 3.**
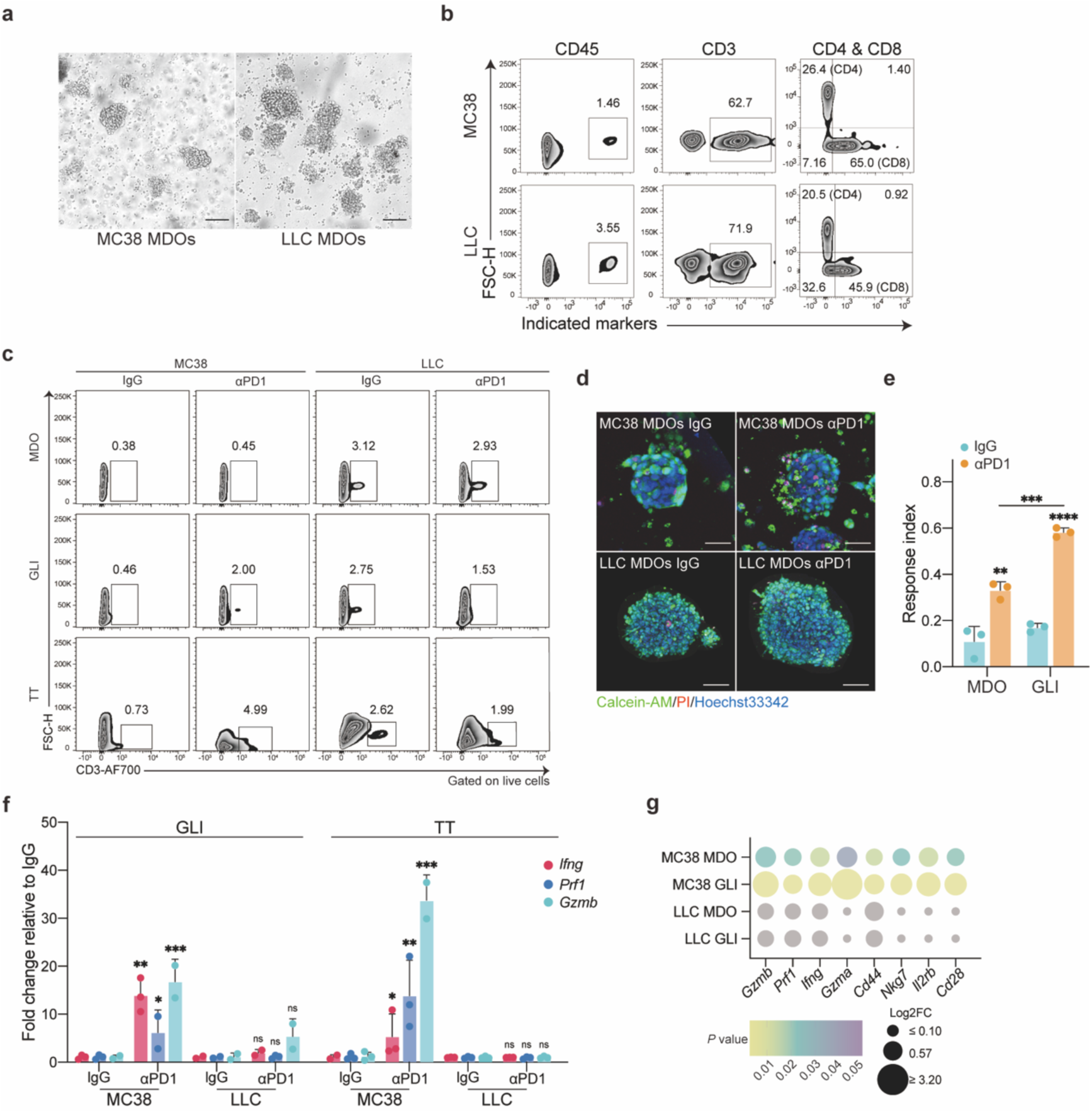
Comparison of the GLI co-cultures and the parental mouse tumors. **a,** Bright field images of MC38 and LLC MDOs. Scale bars: 50 μm. **b,** FACS analysis of the immune components in MC38 and LLC MDOs. **c,** FACS analysis of CD3^+^ T cells in MC38 and LLC MDOs, GLI co-cultures and TTs following IgG or αPD1 treatment, from a single representative experiment with n=3 biological replicates for each tumor line. **d,** Fluorescence microscopy of MC38 and LLC MDOs on day 6 showing the live and dead cells. Scale bars: 50 μm. **e,** Ri of MC38 MDOs mono-culture and GLI co-culture following IgG or αPD1 treatment (n=3). **f,** qRT-PCR analysis of CD3^+^ T cells sorted from (**c**) after αPD1 treatment (normalized to control IgG) (n=3). **g,** Dot plot illustrating the fold change of genes associated with T cell activation or toxicity in MC38-derived MDO and GLI models, LLC-derived MDO and GLI models after αPD1 treatment relative to control IgG. The gray in dots indicates that the *P* value is out of range. (\**P*<0.05, \*\**P*<0.01, \*\*\**P*<0.001, \*\*\*\**P*<0.0001, ns: no significance by unpaired student’s *t* tests)

**Extended Data Fig. 4.**
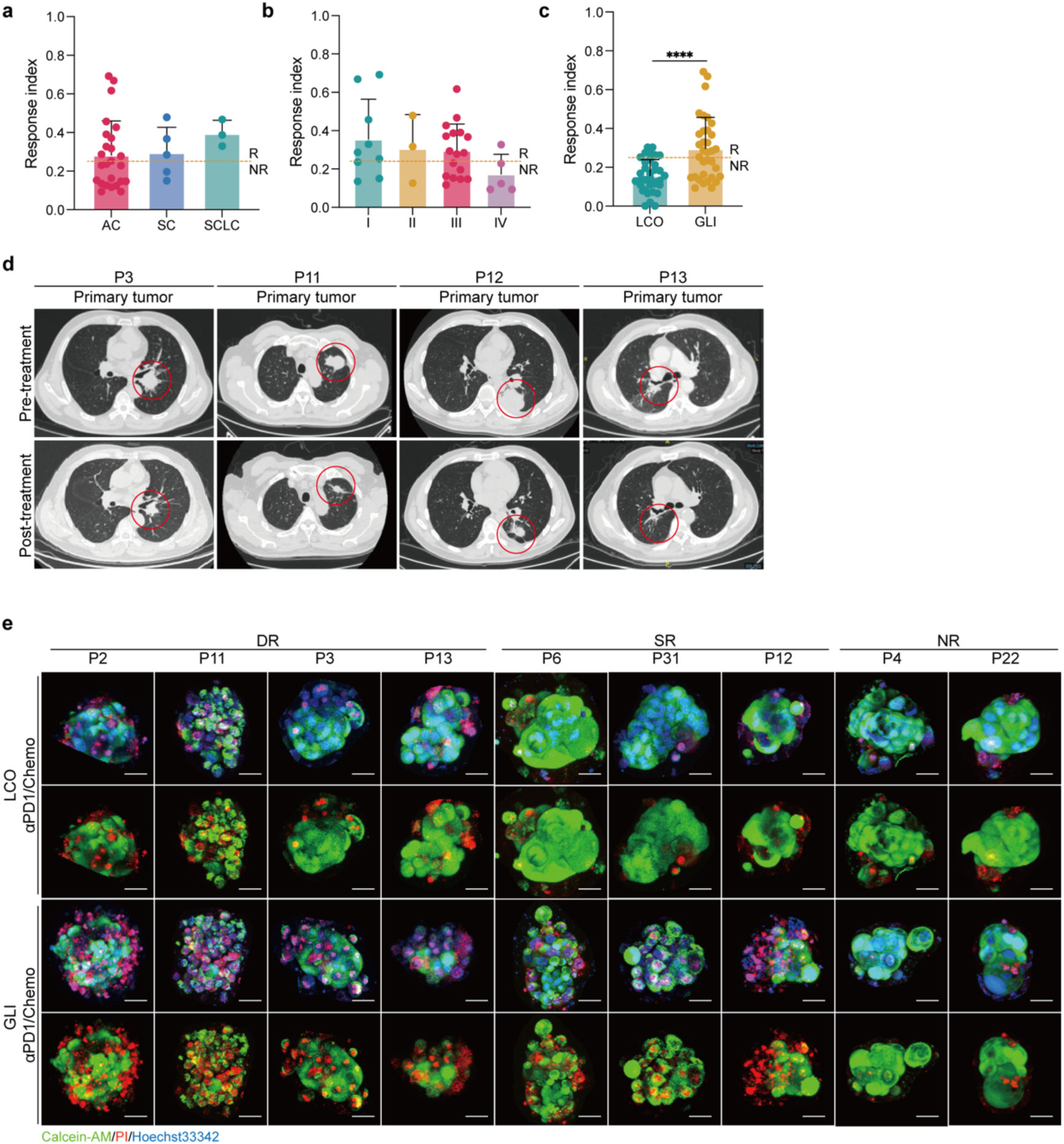
Response of patients and the corresponding *ex vivo* models to immunotherapy. **a,b,** Bar plots comparing the Ri of GLI models derived from lung cancer patients with different subtypes (**a**) and stages (**b**) treated with αPD1. **c,** Bar plots comparing Ri of LCO and GLI models from all patients following αPD1 treatment (\*\*\*\**P*<0.0001 by paired student’s *t* tests). **d,** CT scan images showing the change of lesions in P3, P11, P12 and P13 pre- and post-immunotherapy. The red circles indicate the primary tumor. **e,** Fluorescence microscopy of LCO mono- and GLI co-cultures after αPD1/Chemo treatment from 9 patients. The first line and third lines displayed LCOs staining with Calcein-AM (green), PI (red) and Hoechst 33342 (blue), the second and fourth lines displayed LCOs staining with Calcein-AM (green) and PI (red). Scale bars: 50 μm.

**Extended Data Fig. 5.**
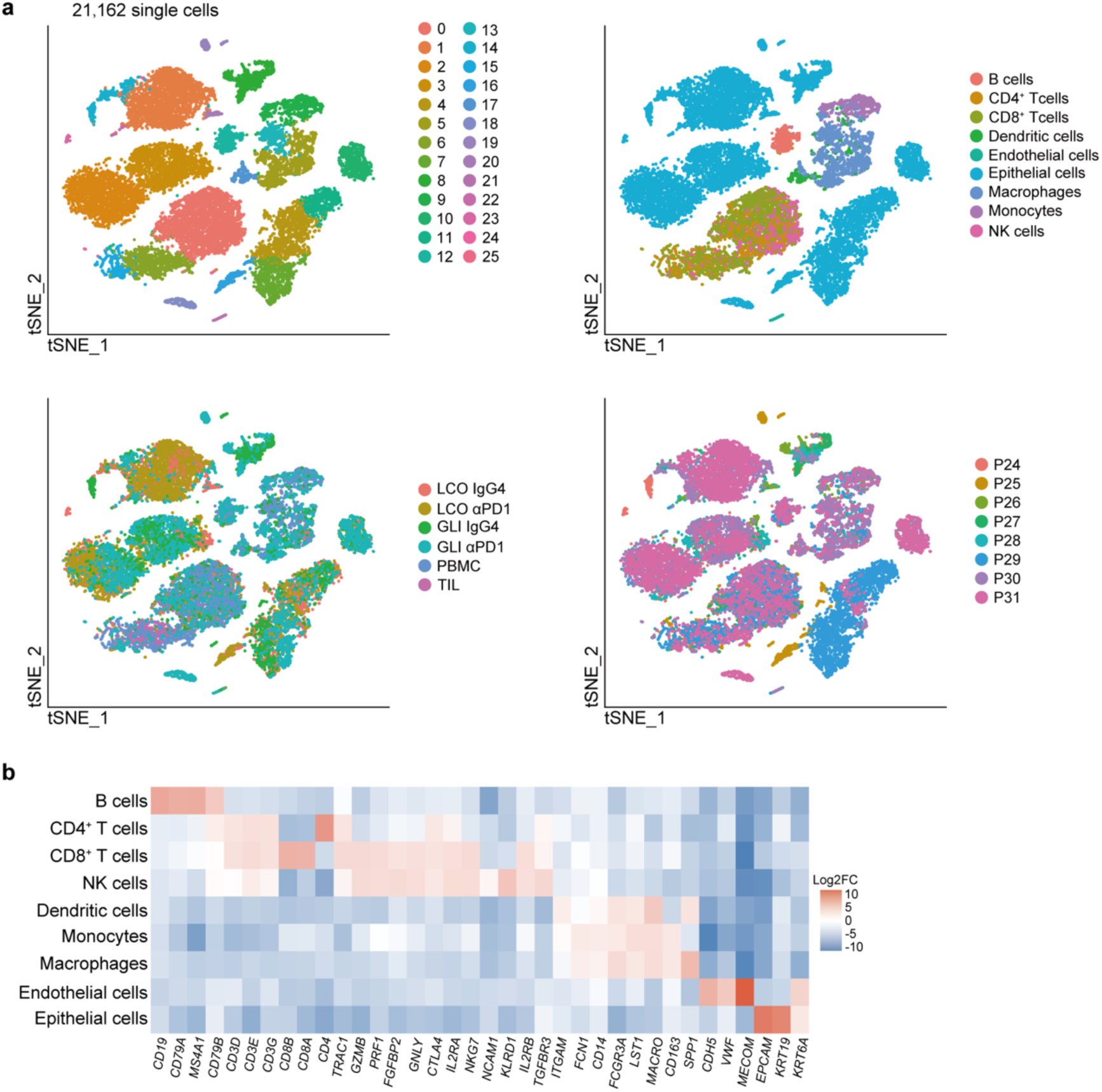
Annotation of the scRNA-seq data of the 8 lung cancer samples in this study. **a,** t-SNE visualization plots of 21,162 cells derived from 8 lung cancer samples labeled by unsupervised clusters (top left), cell types (top right), culture conditions (bottom left) and patient ID (bottom right). **b,** Heatmap showing average expression levels of marker genes associated with the annotated cell types.

**Extended Data Fig. 6.**
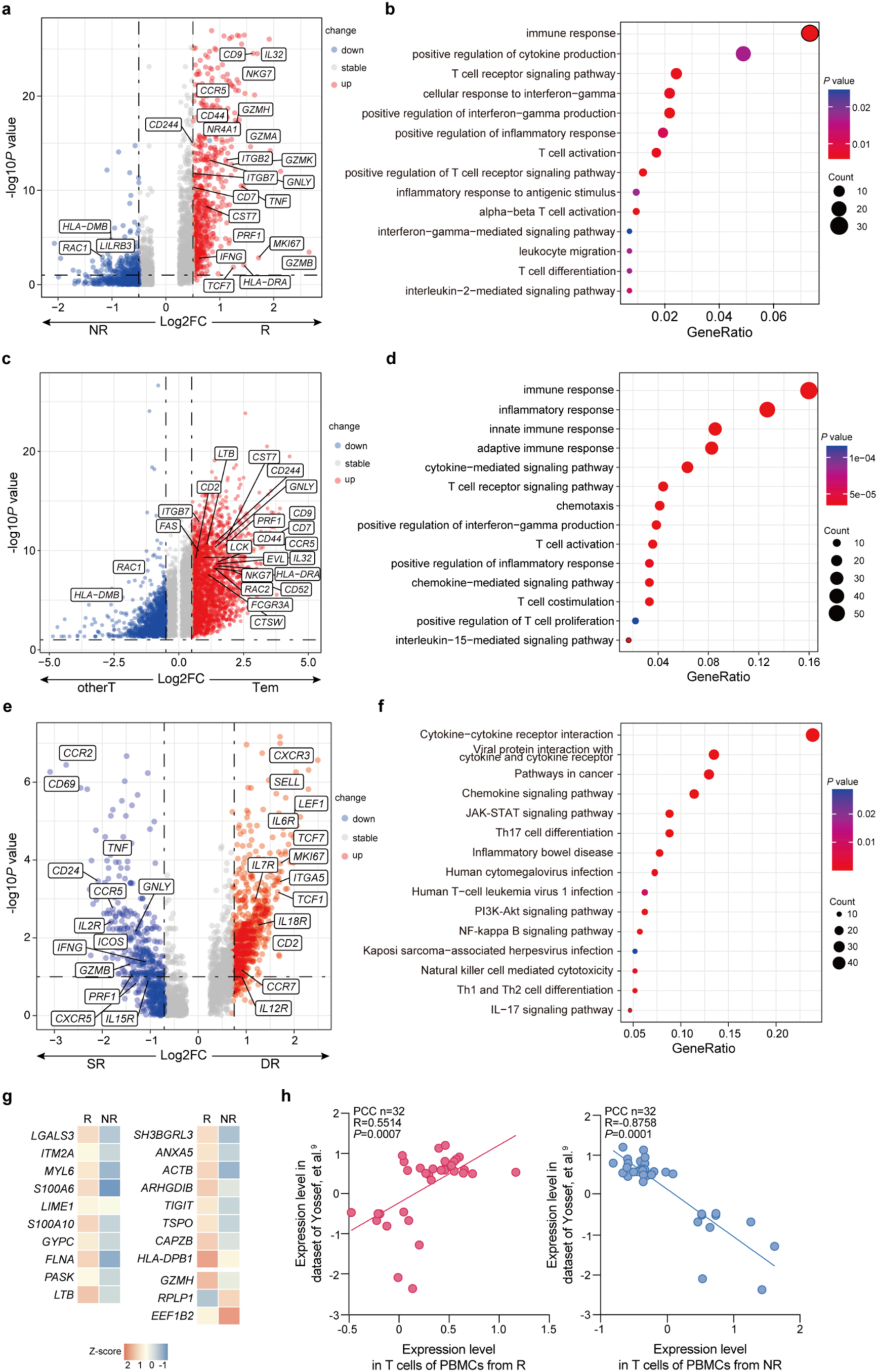
Bioinformatic analysis of T cells in PBMCs. **a,** Volcano plot showing DEGs of PBMC-Ts from the R group relative to the NR group. Each dot denotes an individual gene. **b,** GO enrichment analysis of the DEGs in (**a**). **c,** Volcano plot showing DEGs of Tem relative to other T cells in PBMCs. Each dot denotes an individual gene. **d,** GO enrichment analysis of the DEGs in (**c**). **e,** Volcano plot showing DEGs of PBMC-Ts from the DR relative to the SR group. **f,** KEGG enrichment analysis of the DEGs in (**e**). **g,** Heatmap showing expression levels of genes related to neoantigen-specificity^9^ in PBMC-Ts from the R and NR groups. **h,** PCC analysis of the expression levels of genes related to neoantigen-specificity between our data and the reference^9^ dataset. Each dot represents a gene. Note the positive correlation of the R group (left) and the negative correlation of the NR group (right).

**Extended Data Fig. 7.**
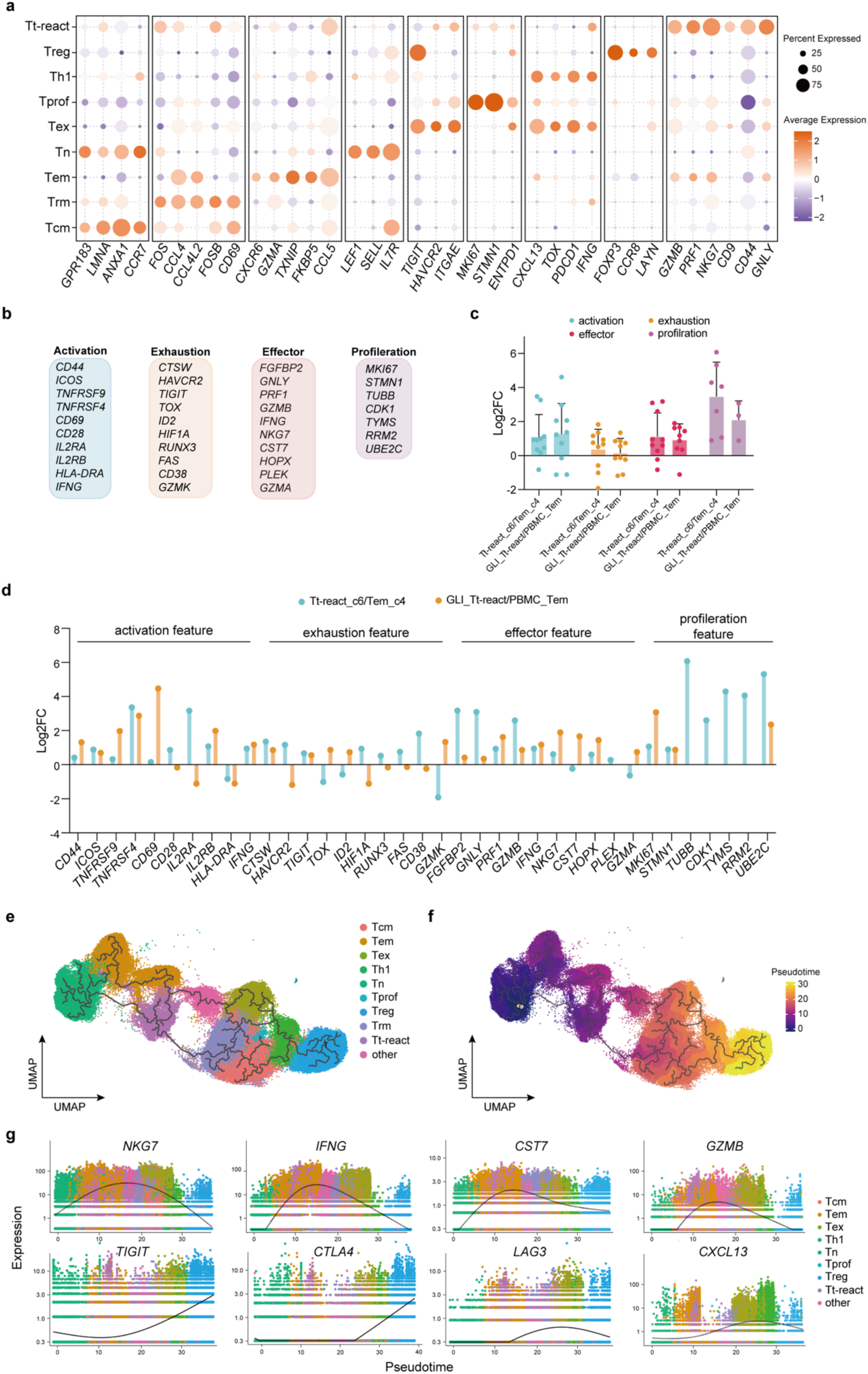
Transcriptome features of Tt-react. **a,** Dot plot displaying the expression levels of celltype-specific genes across 9 distinct T cell subclusters. **b,** Representative genes associated with activation, exhaustion, effector and proliferation as reported.^38,62,63^ **c,d,** Bar plots showing the fold change of expression levels for the genes in (**b**) from Tem_c4 (c4 in fig **4b**) to Tt-react_c6 (c6 in fig **4b**) and Tem in PBMCs to Tt-react in GLI co-cultures. **e,f,** UMAP visualization plot illustrating development trajectory inferred by Monocle3, with cells colored according to cell types (**e**) and pseudotime (**f**). **(g)** Analysis of gene expression patterns along the inferred trajectory, colored by cell types.

**Extended Data Fig. 8.**
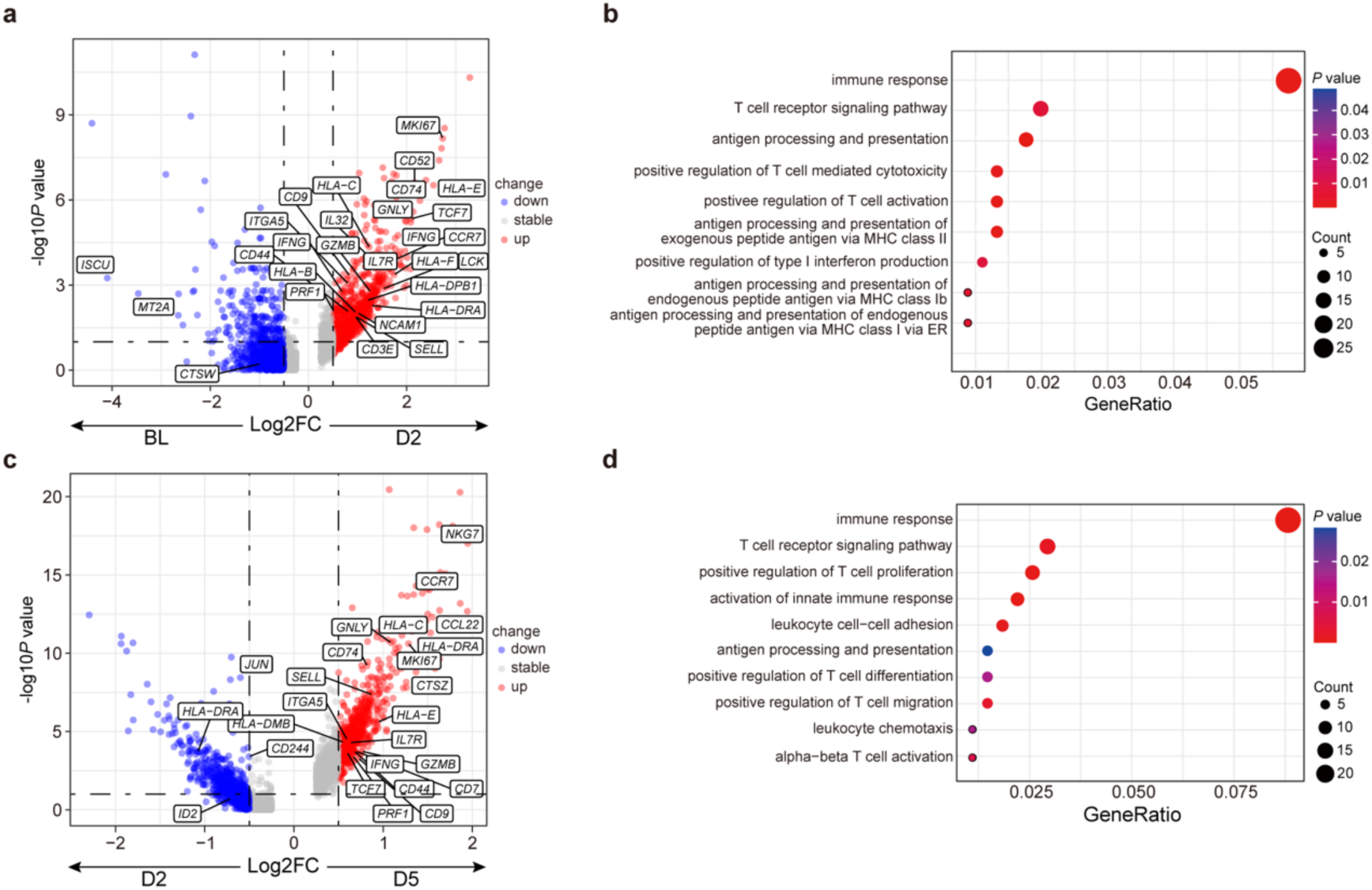
Analysis of T cell dynamic change when encountering LCOs. **a,c,** Volcano plots showing the up- and down-regulated genes of T cells in the P29 GLI models from BL to D2 treated with αPD1 (**a**) and from D2 to D5 treated with αPD1 (**c**). Each dot denotes an individual gene. **b,d,** GO enrichment analysis of the DEGs in (**a**) and (**c**), respectively.

**Extended Data Fig. 9.**
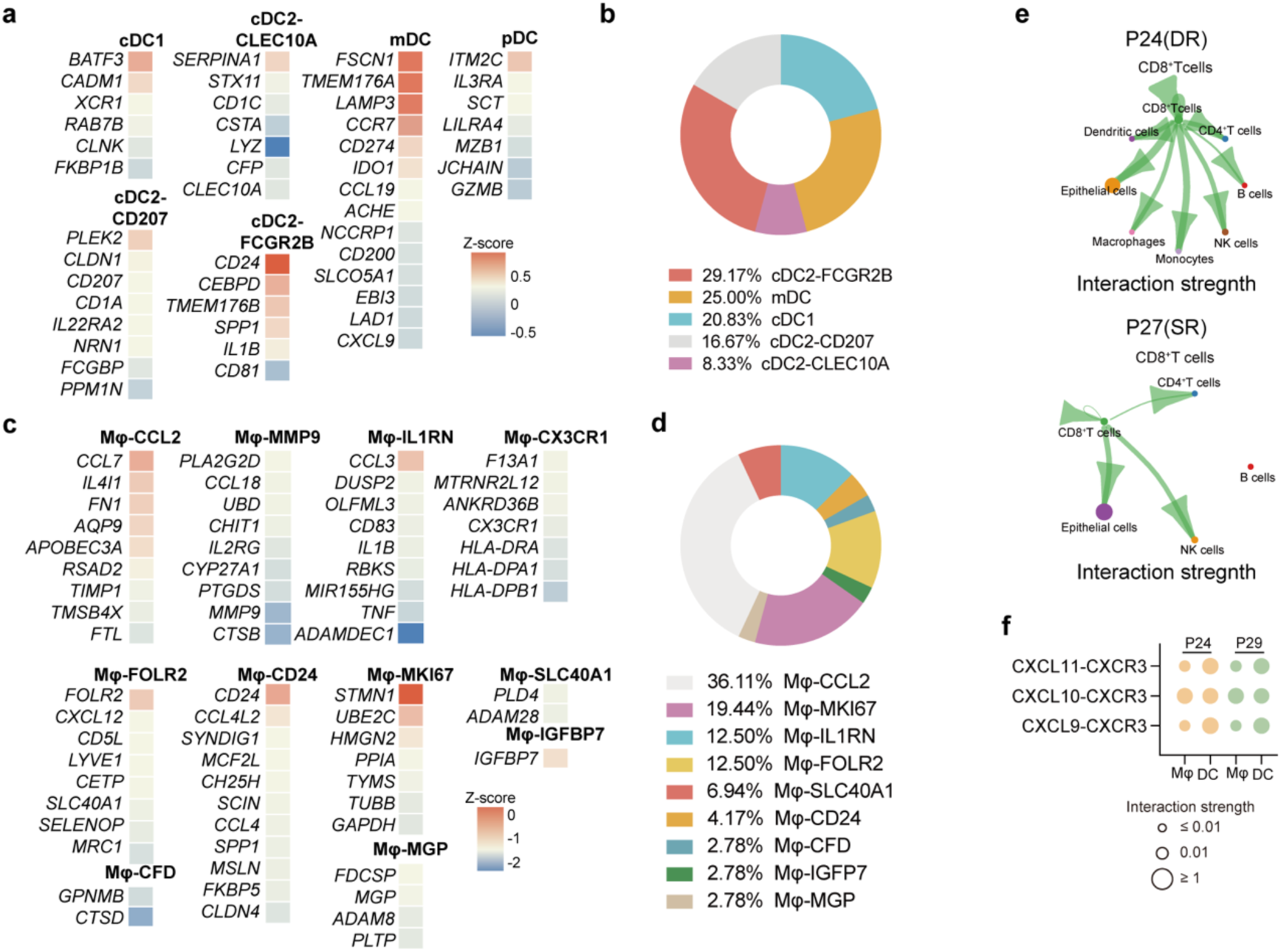
Subtypes of DC and Mφ in LCOs. **a,** Expression levels of genes related to different subtypes of DCs in LCO models from the DR group.^13^ **b,** Proportions of different subtypes of DCs in the DR group. **c,** Expression levels of genes related to different subtypes of Mφ in LCO models from the DR group.^13^ **d,** Proportions of different subtypes of Mφ in the DR group. **e,** Circular plots of interaction strength in intercellular communication between CD8^+^ T cells and other cell types in P24 (DR) and P27 (SR). **f,** Interaction strength of CXCL9/10/11-CXCR3 between T cells and Mφ or DCs in P24 (DR) and P29 (DR).

